# *In Silico* Predictive Modeling of CRISPR/Cas9 guide efficiency

**DOI:** 10.1101/021568

**Authors:** Nicolo Fusi, Ian Smith, John Doench, Jennifer Listgarten

## Abstract

The CRISPR/Cas9 system provides unprecedented genome editing capabilities; however, several facets of this system are under investigation for further characterization and optimization, including the choice of guide RNA that directs Cas9 to target DNA. In particular, given that one would like to target the protein-coding region of a gene, hundreds of guides satisfy the basic constraints of the CRISPR/Cas9 Protospacer Adjacent Motif sequence (PAM); however, not all of these guides actually generate gene knockouts with equal efficiency. Leveraging a broad set of experimental measurements of guide knockout efficiency, we introduce a state-of-the art *in silico* modeling approach to identify guides that will lead to more effective gene knockout. We first investigated which guide and gene features are critical for prediction (*e.g.*, single- and di-nucleotide identity of the gene target), which are helpful (*e.g*., thermodynamics), and which are predictive but redundant (*e.g*., microhomology). We also investigated evaluation measures for comparing predictive models in the present context, suggesting that Area Under the Receiver Operating Curve is not ideal. Finally, we explored a variety of different model classes and found that use of gradient-boosted regression trees produced the best predictive performance. Pointers to our open-source software, code, and prediction server will be available at http://research.microsoft.com/en-us/projects/azimuth.

## INTRODUCTION

One of the most exciting new developments in molecular biology is the CRISPR/Cas9 genome-editing technology^1^. CRISPR (Clustered Regularly Interspaced Short Palindromic Repeats) and their associated endonuclease genes (*e.g.*, Cas9) have been implicated in adaptive immunity in bacteria and archaea; in particular, the CRISPR/Cas9 system is known to function in these organisms by cutting foreign DNA that attempts to invade the host. However, by transferring components of this naturally-occurring defense mechanism into any cell and organism of choice, researchers have developed the ability to precisely edit the genome on a cell-by-cell basis, in any organism. Such precise, reliable and generic genome-editing capability has long been a goal of molecular biology, both as a means to understand gene function and, ultimately, to enable precision medicine through corrections of individual-specific mutations in genes. In contrast to earlier technologies with poor efficiency, cumbersome target design and difficulty in achieving multiple mutations^2^, CRISPR/Cas9 holds far greater promise—it has already been used to perform genome-wide knock-out screens to find genes that provide resistance to drugs or are implicated in disease^3–5^, and to explore the modification of various cells in models of HIV^6^, cystic fibrosis^7^, autism^8^, and muscular dystrophy^9^.

Briefly, the CRISPR/Cas9 system for gene editing/knockout is implemented as follows. The user designs a gene-specific single guide RNA (which we will refer to simply as a guide throughout)—a 20 nucleotide string that is complementary to a portion of the target gene and which must lie adjacent to the Cas-specific Protospacer Adjacent Motif (PAM), which for Cas9, from *S. pyogenes,* is any nucleotide followed by two Gs—that is, the motif “NGG”^10^, where N is a wildcard nucleotide. The guide and Cas9 are then introduced into the cell, for example through the use of lentiviral vectors^11^, where the guide/Cas9 complex then gets recruited to the target gene sequence through basepairing between the guide and the gene sequence itself. At this point, Cas9 cuts both strands of DNA, which then get repaired by endogenous pathways. Depending on whether the goal is editing of a gene sequence, or a knockout edit, one of two repair mechanisms is induced. For gene editing, homology-directed repair incorporates a user-provided template that specifies desired edits to the target region; this tends to be a low-efficiency process. In contrast, gene knockout relies solely on the non-homologous end joining (NHEJ) repair mechanism, an inherently faulty processes that frequently introduces insertions or deletions (indels) to cause frameshifts and potentially create null alleles; many of the faulty repairs have the effect of knocking out the function of the gene, fulfilling the intended goal. Both of these repair processes depend on properties of both the guide and gene, such as whether the target DNA is accessible for base-pairing with the guide, thermodynamic and other properties of the guide, as well as sequence properties in the nearby area of the gene (beyond the 20 nucleotide matching portion). Furthermore, for gene knockout, even if a mutation is created by virtue of the faulty repair, generation of a loss-of-function allele is likely to depend on where the double-stranded break occurs within the protein-coding region (*e.g.*, if the guide is targeting the gene nearer the N’ or C’ terminus), and also whether the indel disrupts the reading frame. Factors influencing these outcomes are starting to be investigated^11^ but are not yet fully understood.

CRISPR/Cas9 technology is still in its infancy, with many important challenges remaining. A case in point is the selection of an RNA guide that enables the system to identify its target location in the genome. While hundreds of possible guides may be available in principle for a given task, not all of them are effective in practice^11^. Moreover, it is not feasible to routinely test all possible guides for a target gene to see which ones work, highlighting the need for computational methods to bridge the gap. Therefore, in the present work, we build predictive models to identify high activity and low activity guides for any gene, using the principles of statistical estimation both to choose a suitable model and to tune its parameters. In doing so, we have enabled more rational and effective use of the CRISPR system. Our corresponding prediction software and server will allow the broader biological community to accurately identify the best guide for their genes of interest, and also to easily build on our insights and algorithms as more data becomes available.

The present work is focused on modeling of on-target effects, that is, how likely a guide is to cut intended DNA targets. We do not model off-target effects, whereby guide RNAs act unintentionally at imperfectly-matched sites. Currently there is a paucity of data for modeling off-target effects, but we plan in future work to combine these two aspects of the problem. Also note that the impact of off-target effects depends partly on the intended experimental aims^12^—for personalized medicine, for example, off-target effects could be deadly; however, for screening assays, such as those examining the functional consequence of gene knockout, off-target effects can be mitigated by combining results obtained from several differehnt guide sequences that share a common target. In addition to examining on-target effects, we also focus on gene knockout with the CRISPR/Cas9 system, rather than editing, doing so with human and mouse genes. We believe that the data-driven models we derive capture information about guide efficacy that should be relevant, if not necessarily immediately transferrable to other organisms.

There have recently been a handful of studies that attempt to provide CRISPR guide selection by way of some form of model. The CCTop algorithm^13^ creates off-target scores based on a simple function of distance to the PAM, and then combines these to obtain a target site score, which is then combined with distance to the PAM and other factors to yield a final score. Similarly, CRISPR Design^10^, E-CRISP^14^ and CHOPCHOP^15^ also use scoring functions with a handful of parameters determined *a priori*, rather than through a data-driven modeling approach based on principles of statistical estimation, such as we perform herein. To our knowledge there have been only three such data-driven statistical approaches previously taken, including our own. All are based on either Support Vector machines (SVM)^4^, logistic regression^16^, or both^11^, and all using classification rather than regression as we use herein. In our previous work, the SVM was used to perform feature selection, followed by a logistic regression model trained to make a binary classification of the top 20% most effective guides versus the bottom 80% effective guides^11^. In other work, either the SVM or logistic regression were used directly, implicitly performing feature selection^4,16^. These predictive modeling approaches to CRISPR guide design can be substantially improved upon in several ways-- the main goal of this manuscript. Additionally, we elucidate broad types of features that are useful for guide prediction, and expose particular features within broad classes of features, such as which dinucleotides at what position are most likely to be associated with an efficient guide rather than a less efficient one.

There is a body of related literature concerning computational approaches for designing small interfering RNA (siRNAs)^17,18^, wherein properties of siRNA sequences are encoded into features for predicting efficiency, including sequence-based features such as GC content, melting temperature of the siRNA, and secondary structures of the target mRNAs. These features were used in conjunction with different models, such as Decision Trees or Neural Networks, to find good predictive models.^17–19^ Although inspiration can be drawn from these computational approaches, CRISPR/Cas9 guide design is an inherently different molecular mechanism, with different amount and types of data available, and therefore requiring independent investigation of computational approaches.

## RESULTS

### Overview

In our earlier work^11^, two fundamental assumptions were made which, when relaxed, substantially increase the predictive performance. The first was to assume that exactly 20% of the guides for each gene were successful at generating gene knockout, and that the remaining 80% were not. As a consequence of that modeling decision, we previously trained a two-class *classification model* (successful guide, vs. not successful). However, we later hypothesized that discretizing the measured activity of the guides, so dramatically, was likely to obliterate important information for prediction. Indeed, as we will demonstrate, this turned out to be the case. Consequently, in the present work, we instead retain a real-value for each guide (the normalized rank^11^—the ranks of the scores for each gene-guide pair, normalized to lie between 0 and 1) and then trained a *regression model* to predict these scores. The second assumption made in ref. ^11^, inherent in logistic regression, was that the (log of the ratio of the probability of success to) probability of failure of knockout for a guide is a linear combination of the features, rather than explicitly comprising more complex interactions among the features. Thus, the logistic regression approach precluded explicit interaction among predictive features such as “if the guide has property X and property Y at the same time then it is more likely to be successful.” Therefore, in the present work, we systematically examined the performance of a variety of models, finding that a class of models which allow for a much richer interaction of features in an explicit manner—regression trees^20^—enabled improved predictive performance.

While modeling choices are critically important to achieving the best predictions possible, the criterion for evaluation of models is equally important. Without a good evaluation criterion, it is impossible to choose the best model or features to use going forward. In refs. ^4,11,16^, the criterion of evaluation used was the Area Under the Curve of the Receiver Operating Characteristic (AUC-ROC or simply AUC). However, as we shall see, in moving from a classification to a regression approach, we find that binarization of the data is lossy when evaluated by the AUC, and thus an evaluation criterion predicated on such a binarization (AUC) is also lossy. Rather, one should use an evaluation criterion that uses the real-valued normalized ranks, such as the mean-squared error, the out-of-sample predictive log likelihood, or various forms of correlation between the predictions and actual normalized ranks. As we discuss in more detail later, we opted to use the Spearman correlation of all guide predictions with the score ranks derived from laboratory experiments. Finally, aside from modeling and evaluation considerations, the actual features of the guide used in the models can dramatically alter the predictive performance. We here augment the features used in ref. ^11^ to include thermodynamic properties of the target region and other features of the guide, showing that together they lead to a statistically significantly better performing model.

### Experimental Setup

We used two primary data sets, which we will refer to as “FC” and “RES” because the methods used to detect successful knockdowns were flow cytometry and resistance assays, respectively. The FC data^11^ consisted of 1,831 guides targeting three human (CD13, CD15, CD33) and six mouse (Cd5, Cd28, H2-K, Cd45, Thy1, Cd43) genes, all producing cell-surface markers which could be assayed by flow cytometry. (There were 1,841 guides to start, however, 10 were removed due to ambiguous mapping to the reference genome). The FC knockout efficiency was evaluated by sorting cells based on whether the protein coded for by that gene was present or not, and then counting relative abundances—a very direct measure of gene knockout.^11^ The RES data (Doench *et al.,* in submission) comprised a further 2,549 unique guides targeting 8 additional human genes (CCDC101, MED12, TADA2B, TADA1, HPRT, CUL3, NF1, NF2), whose knockout success was inferred from resistance to one of three drugs (vemurafenib, 6-thioguanine, selumetinib): if CRISPR/Cas9 with the guide successfully knocked out one of the genes of interest, then those cells would be able to survive exposure to the drug (for certain drug-gene pairs). For both the FC and RES data, guide “scores” representing how successful a particular guide was at knocking out its target gene were computed: the guides for each gene are assigned a score between 0 and 1, which is a normalized rank derived from the change in abundance after cell sorting/drug exposure (see the Methods section for more details and rationale).

For many of our experiments we conducted comparisons across three versions of our data: (i) on the FC data alone (9 genes across 1,831 guides), (ii) on the RES data alone (8 genes, 2,549 guides), (iii) on the combined FC and RES data (17 genes, 4,380 guides). In all cases unless otherwise noted, we did leave-one-gene out, so as to best understand the generalization ability of our models for genes never seen before. Unless noted otherwise, the guide features were the same as those used in ref. ^11^, which consisted of position-specific sequence features for individual (order 1) and pairwise (order 2) nucleotides in the 30mer sequence (the 20mer guide plus context on either side) along with GC counts for each guide (*e.g.*, Supplementary Figure 4). Further details, including statistical tests used to compare performance, are explained in the Methods section.

### Classification, Regression and Evaluation

We first investigated whether the use of the real-valued guide scores could yield better predictive results than using the 20%/80% binary discretization used in ref. ^11^ To make the comparison as fair as possible, we first retained the same evaluation criterion as ref. ^11^, the AUC. As seen in Figure 1a, L1-regularized linear regression (a regression model) systematically outperforms L1-regularized logistic regression (a classification model), when trained on the same data with the same features, using the AUC criterion. This result demonstrates that the real-valued scores clearly are more informative than the 20%/80% discretized data.

**Figure 1.**
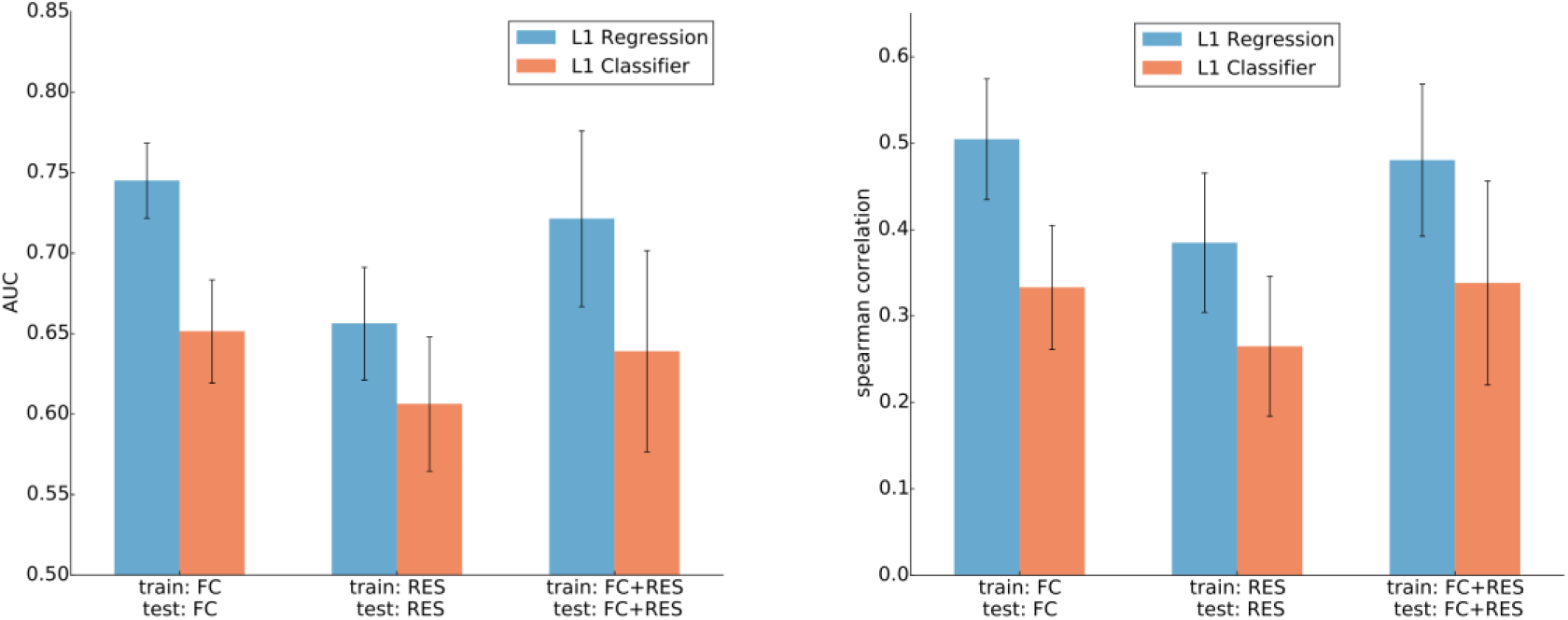
Comparison of two comparable predictive models, one of which uses only a 0/1 binarization of the guide scores (L1 Classifier), and the other that uses the ranks derived from guide scores (L1 Regression). Left figure uses the AUC, while the right uses Spearman correlation. For all three data sets, and across both criteria, the regression approach outperforms the classification approach. Each bar shows the mean value across all genes with the error bars denoting one standard deviation across the genes. Statistical significance of the improvement in Spearman correlation when using regression over classification is, for each of FC, RES and FC+RES data, respectively p<1 × 10^-16^, p=5.4× 10^-13^ p<1 ×10^–16^.

Thus, we decided to use a performance criterion that could make use of these real-valued scores. For practical purposes, what we care most about in the present problem setting is correctly ranking, say, the top 5 guides for each gene, and ignoring the ranking of the other guides. The information retrieval (IR) community has directly analogous problems, such as providing the top 10 web pages to a search query while ignoring all other possible web pages. To solve this problem, the IR community uses evaluation criteria such as normalized discounted cumulative gain^21^ (NDCG), in which the top of the list gets more weight in the metric than the bottom, and that at some point, the rest of the list is completely ignored in the metric. However, when we attempted to adopt this measure for the CRISPR prediction problem, we discovered that any performance criterion that discards or discounts information from all but the top-ranked items requires many, many “queries”, or in our case genes, to yield low enough variance on the criterion such that it is useful. In our case, we had at most 17 genes (when combining all available data), and the resulting NDCG variance seemed too high to be useful. We found that the Spearman correlation of all predictions with normalized ranks to be a more suitable criterion. In contrast to AUC, our Spearman correlation makes use of important rank information, and, does not require that each gene have precisely 20% success among all attempted guides. Using this Spearman correlation criterion, we again saw that the regression model systematically outperformed the classification model (Figure 1b). As a consequence of the results in this section, going forward, we make use of only the Spearman correlation, and only regression models rather than classification models.

For completeness, we also compared the performance of the three previously-used classification modeling approaches and variations of these: SVM using order 1 nucleotide features^4^, SVM using up to order 2 nucleotide features followed by logistic regression^11^, and logistic regression using order 1 nucleotide features^16^. We also augmented these models to use up to order 2 nucleotides when they had not already done so. When assessed on the FC and RES datasets, we observed the best performance from SVM plus logistic regression. We also observed that when the SVM or L1 logistic regression approaches are augmented by order 2 features, they still performed worse than the SVM plus logistic regression from ref. ^11^ on these datasets (Supplementary Figure 1).

### Additional features

So far, we have used the original features proposed in ref. ^11^, which comprise: (a) all single position-specific “one-hot” encoded nucleotide features (*e.g.*, is there a T in position 3 of the 30mer, described in more detail in Methods), (b) all two-nucleotide-adjacent position-specific “one-hot” encoded nucleotide features (*e.g.*, is there a TG in positions 3 and 4 of the 30mer), (c) the GC content/count—how many times G and C appear in the 20mer guide, and whether that count is greater than or less than 10. However, we hypothesized that further features could improve the prediction. In particular, biochemical and structural studies have shown that particular parts of the 20mer guide participate in step-wise association with DNA^22,23^, suggesting that the thermodynamic properties of different guide segments may yield useful features.

Altogether, the additional features were: position-*independent* nucleotide-features (*e.g*., how many “A”s, or how many “AT”s, irrespective of where in the 30mer); position of the guide within the gene (amino acid cut position; percent peptide; percent peptide<50%); “NGGN” interaction (the identities of the nucleotides (nts) on either side of the invariant GG of the PAM); melting temperature of (i) the 30mer (20mer guide plus context), (ii) the 5mer immediately proximal to the NGG PAM—positions 16-20 of the 20mer guide, (iii) the 8mer in position 8-15 in the 20mer guide, (iv) the 5mer in positions 3-7 of the 20mer guide. These features improved the model in a statistically significant manner for all three data sets (Figure 2). We also tried including the gene length; the gene GC content; the gene melting temperature; and the gene molecular weight, but found at most a modest improvement and so did not retain these features.

**Figure 2.**
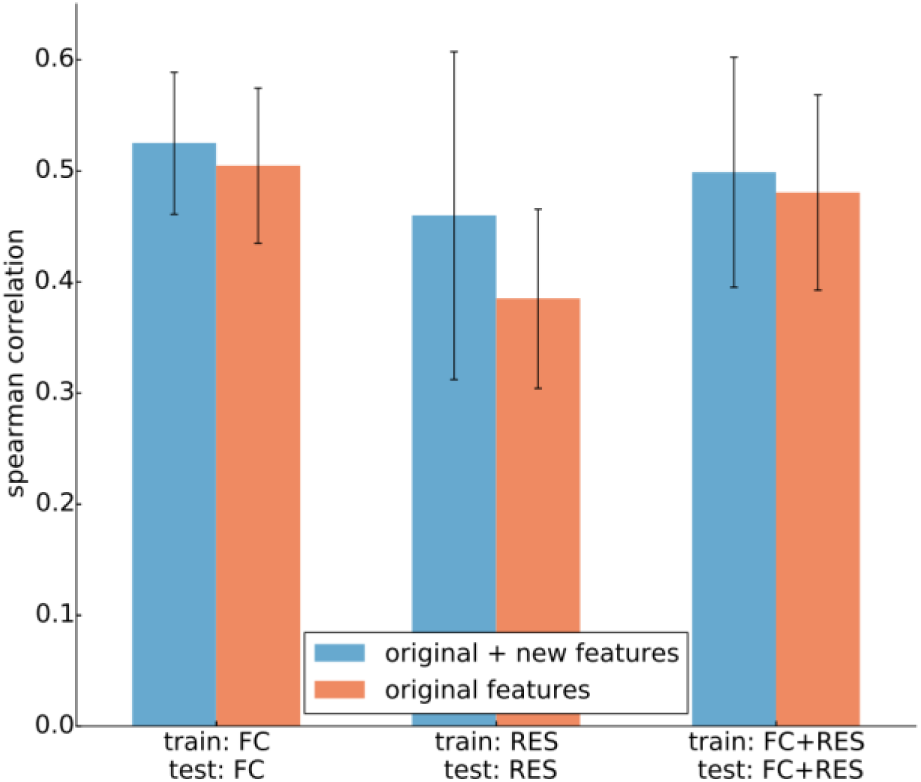
Spearman correlation performance of L1-regularized linear regression when using the original features from ref. ^11^ compared to adding in new features as described in the text. Each bar shows the mean value across all genes with the error bars denoting one standard deviation across the genes. The statistical significance of the improvement in Spearman correlation when using new features for each of FC, RES and FC+RES data was respectively p=4.2 × 10^-3^, p<1 × 10^-16^, p=2.32 × 10^-4^.

Finally, we tried adding microhomology features^24^. These features did not yield improved predictive performance, but were on their own, predictive. In particular, we took the 30mer guide and context sequence already being used and augmented a further 9 nucleotides to the left and 21 to the right to yield a 60mer centered on the cut site. Then we used the method from ref. ^24^ to produce a microhomology score as well as an out-of-frame score, and added these as features to our model. On the FC data, the L1 regression benefited slightly (median Spearman correlation across genes of 0.523 vs. 0.533) from the use of these features, but once we settled on a final model class, described later, we found that these features no longer helped (median Spearman correlation across genes of 0.526 in both cases), presumably because the richer models can make better use of the other features, making these redundant.

### Systematic comparison of different predictive models

Having established that regression models with additional features were the best way forward, we next systematically investigated the predictive performance of five different regression models, and also compared these to the best previous classification approach.^11^ The five regression models were: (i) Gradient-boosted regression tree, (ii) L1-regularized linear regression, (iii) L2-regularized regression, (iv) linear regression, and (v) random forest regression. These models differ on several axes. One axis is whether or not the features are combined only linearly (L1-regularized regression, L2-regularized regression, linear regression), or can be combined in highly non-linear ways (random forest regression, Gradient-boosted regression tree). Although linearity works well in many settings, flexibility offered by non-linear models can sometimes yield substantially better results. Another axis is how the model mitigates over-fitting, which occurs when there is too much freedom in the model relative to the amount of data available to fit the model parameters. Linear regression does nothing to mitigate the problem, while L2-regularization shrinks the weights to zero according to a squared penalty on the weights, and L1-regularization induces a sparse set of weights (only some of them are non-zero). Gradient-boosted regression trees and random forests prevent over-fitting by limiting the depth of their trees. A third axis that differs substantially among these models is whether it is an “ensemble” method—that is, a method that actually uses many (*e.g.*, hundreds) of some base model (*e.g.,* regression tree) and combines them together in some data-driven manner. Both Gradient-boosted regression trees and the random forest are ensemble methods. Finally, one practical way in which the models differ is in their need (or lack thereof) to use cross-validation (CV) or some other means to fit the regularization parameter. Both L1 and L2 do require such an approach. While one can use CV to tailor the Gradient-boosted regression tree and random forest settings, we did not here do so, instead using the default settings.

In all but one case, the Gradient-boosted regression tree had statistically significantly better Spearman correlation than all the other methods (Figure 3). Therefore, we concluded that the Gradient-boosted regression tree was the best model among those compared here. (Only on the FC data, compared to L2- and L1-regression, did the Gradient-boosted regression tree fail to yield statistically significantly better performance.)

**Figure 3.**
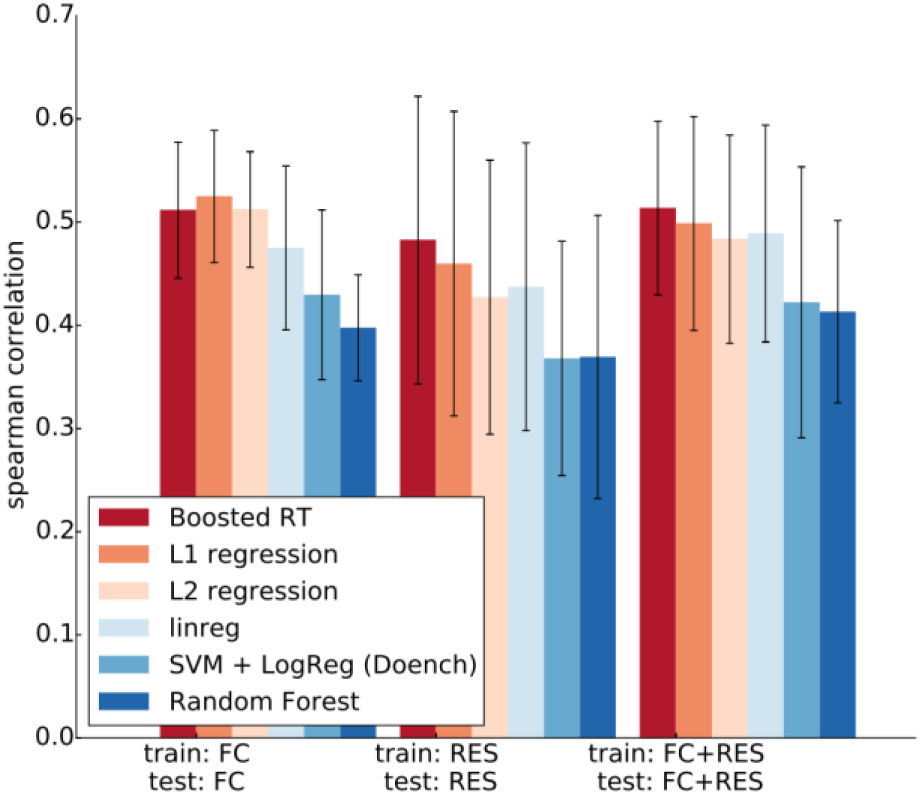
Comparison of the predictive performance of different regression models. “SVM + LogReg (Doench)” denotes the classification approach used in ref. ^11^ Each bar shows the mean value across all genes, with the error bars denoting one standard deviation across the genes. The statistical significance of the improvement in Spearman correlation when using Boosted RT (boosted regression trees) over the other models was, respectively for the FC, RES, FC+RES data, p=5.4 × 10^-2^, *p* = 4.9 × 10^-4^, *p* = 5.3 × 10^-5^ for L1 regression, *p* = **4.5** × **10**^**–1**^, *p* = 2.8 × 10^-15^, *p* = 3.1 × 10^-8^ for L2 regression, p=1.3 × 10^-7^p< 1 × 10^-16^, *p* < 1 × 10^-16^ for linreg, SVM + LogReg, and *p*= 1.5 × 10^-10^p< 1 × 10^-16^, *p* < 1 × 10^-16^for Random Forest. All but the bolded values are significant.

Regression trees encode rich interactions by partitioning data based on input features as one “walks down the tree from the root”. That is, each branching enables the effect of that feature to depend on the branching (value) of previous features, thereby yielding an extremely rich expressive language for the model. Additionally, in using not one but a set of regression trees (a so-called ensemble of regression trees), we were able to improve performance further. The set of trees here was determined and combined according to the rules of the gradient-boosted regression tree algorithm,^25^ a particular version of boosting.^26^ Intuitively, boosting works by first training one regression tree on the training data, as though it were the only model. Then, it measures the error of each training example, and re-weights those with poor performance more highly. Using this re-weighted data, it trains a second regression tree. This process continues iteratively until a specified number of trees has been trained on each newly re-weighted version of the training data set. The final boosted regression tree model is then a weighted sum of the predictions from each regression tree, weighted in inverse proportion to the error of each tree. Although the number of trees and other parameters can, in principle, be learned by cross-validation, here we did not do so. Instead we used a gradient-boosted regression tree learning rate of 0.1, a max tree-depth of three, and 100 regression trees.

Note that we also briefly investigated the use of Gaussian Process Regression^27^, Deep Neural Networks classification^28,29^ (DNN), and Elastic Net regression^30^, but were not encouraged by preliminary results, and therefore left exploration of these models to future work. We note, however, that the present setting is not one in which DNNs are known to work well, that is, a setting in which massive amounts of (usually supervised) data are available. It is also interesting that our preliminary results suggested that L1-regression performed at least as well as Elastic Net, a conclusion concordant with the results in ref.^16^, where the regularization parameter from Elastic Net yielded a strict L1-based model (Elastic Net encompasses both L1 and L2 regression, and blends them in a data-driven manner).

For completeness, we also compared our final approach (gradient-boosted regression tree, “Boosted RT”) to the trained (by the authors) model of Xu et al^16^, as provided with their paper (Supplementary Figure 2, top row). We compared approaches both on our datasets (FC and RES), as well as on the combined three datasets used by Xu *et al.* (Wang ribosomal, Wang non-ribosomal, and Koike-Yusa), which we here refer to as the “Viability” data. To ensure that no test cases appeared in the training data, for our approach we used RES to train when testing on FC, and vice-versa, and used FC+RES for training when testing on the Viability dataset. The actual trained model provided by Xu *et al*. was presumably trained on all of the Viability dataset, in which case their model gains a potentially substantial apparent advantage in performance when also tested on the Viability data, as the training and test sets overlap (this will become apparent in the next paragraph). Despite this, we found no statistically significant difference in performance between the Boosted RT model and the Xu *et al*. model when applied to the Viability data (p=0.22 when comparing the Spearman correlations). Further, relative to its performance on the Viability datasets, we observed a decrease in performance of the Xu *et al*. model when tested on the FC or RES datasets, both relative to their performance on their own data as well as relative to the performance of our model (p<10^-16^ for both FC and RES), even our model never used the same data set for training and testing by virtue of splitting up FC and RES. (The FC data was for some reason not reported on by Xu *et al.*)

Because of the hypothesis that the final model provided by Xu *et al.* had been trained on all of the Viability data and therefore had an upwardly biased estimate of performance on that same data, we also re-trained the Xu *et al*. model (L1 classification, order 1 nuc. features) on the Viability data, and used stratified (by gene) cross-validation so that the training and test sets were never overlapping (Supplementary Figure 2 bottom row). We observed a decrease in performance of the modeling approach used by Xu *et al.* relative to using the already-trained model provided by their paper, suggesting that their model had in fact been trained on all of the Viability data. In this setting, Boosted RT outperforms the Xu *et al.* modeling approach in a statistically significant manner (p=7.0 × 10^-10^). We also trained the Boosted RT model on the Viability data, and then tested it on the FC and RES data sets; in both cases we observed significantly better performance (respectively, p=2.1 × 10^-15^, p=1.7 × 10^-12^). These results suggest that the final, trained Boosted RT model provided herein generalizes well across multiple data sets.

### Feature importance

We concluded earlier that our new features yielded statistically significantly better predictive results when using L1 linear regression. The same was true when using Gradient-boosted regression trees, except on the FC data (Spearman correlation with/without new features here was 0.51 and 0.52). We here highlight the overall most important categories of features (Figure 4), as well as the top ranked individual features (Table 1). The final, fully-learned model on all of our data is available with our associated software in programmatic form, and the complete ranked list of individual features is in the Supplementary Information. Features are ranked according to their average Gini importance over all regression trees in the ensemble. The Gini importance here refers to the decrease in mean-squared error (the criterion used to train each regression tree) when that feature is introduced as a node in the tree. This measure has a close, empirical correspondence with the importance of the feature that would be obtained with a permutation test, and can also be viewed as a relative decrease in entropy provided by splitting on that feature^20,31^. Note that this measurement of importance effectively averages out the effect of the other features, since the addition of a feature most often happens after already having split on a different feature. Additionally, this measure of importance does not convey whether having that feature makes a guide better or worse in the model (favored/disfavored in the terminology of ref. ^11^)—again, because such a notion is impossible for regression trees, in which the effect of one feature is dependent on the presence/absence of other features. This lies in direct contrast to linear [logistic] regression, in which the effect [latent liability] is a weighted sum of the features and therefore allows more direct interpretation of each feature alone, including whether it favors/disfavors activity of a guide. Consequently, in this section, we also investigate the feature weights of the L1-regularized regression, the second-best model, so as to obtain some intuition on the directionality of each feature. Note that it is precisely this extra level of richness in the regression trees that allows them to predict better, but which also makes their use of individual features harder to easily summarize in a human-readable manner.

**Figure 4.**
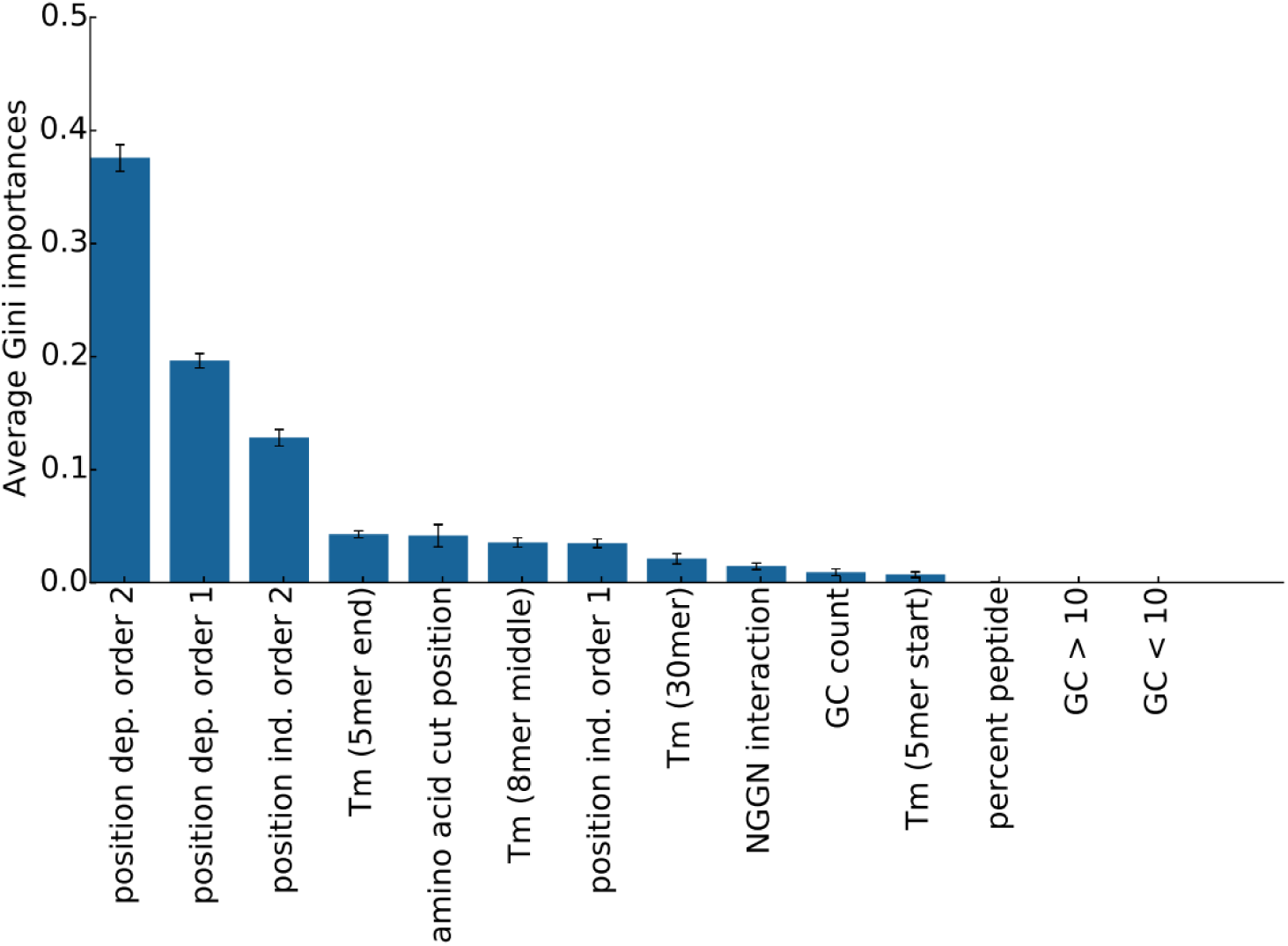
Relative importance of each group of features in the Gradient-boosted regression trees using the Gini importance, for the experiment using the FC+RES data. Similar plots for each of the old and new data are shown in Supplementary Figure 3. “position dep. order 2” denotes position-dependent features for adjacent paired nucleotide features, whereas “position ind. order 2” denotes the same thing, but aggregated over all positions.

**Table 1:** Relative importance of each of the top 10 individual features in the Gradient-boosted regression trees using the Gini importance, for the experiment using the FC+RES data. For single and di-nucleotide position-specific features, the value of nucleotide(s) is reported first in the feature name, then conjoined with a “_”, and then finally with the position. Therefore “AA_18” refers to a dinucleotide features in positions 18 and 19, with value A and A. In contrast, “GG” is a position-independent feature that simply counts the number of times “GG” occurs.

|  | **Feature** | **Importance** | **Std. Dev.** |
| --- | --- | --- | --- |
| 1 | percent peptide | 0.093 | 0.009 |
| 2 | Tm (5mer_end) | 0.043 | 0.003 |
| 3 | amino acid cut position | 0.042 | 0.010 |
| 4 | Tm (8mer_middle) | 0.035 | 0.004 |
| 5 | G_20 | 0.027 | 0.002 |
| 6 | GG | 0.023 | 0.004 |
| 7 | TT_17 | 0.021 | 0.002 |
| 8 | Tm (entire 30mer) | 0.021 | 0.005 |
| 9 | A | 0.019 | 0.002 |
| 10 | AA_18 | 0.016 | 0.002 |

Looking at the broad sets of features in Figure 4, we see that the most important features turn out to be the position-dependent nucleotide features—both the singletons, and the adjacent pairwise ones. Next-most important are the position-independent pairwise nucleotide features, followed by several of the thermodynamic properties of the guide. When we drill down to specific individual features, among the top 10 are positional features in the target gene, the GC count, and the NGGN interaction features (where the feature consisted of which two letters occur in each of the “N”s in NGGN, where NGG is the PAM). It stands to reason that with further investigation, one might find even more features that yield increased performance.

Note that the ranking of individual features listed in Table 1 may seem incongruous with that in Figure 4, but this is simply because Figure 4 is ordering based on the average importance of all features within a specified group, whereas Table 1 is ordering based on every individual feature regardless of group. Note that both “amino acid cut position” and “percept peptide” are features that indicate the position of the target within the gene, and both are important predictive features. The latter is restricted to lie in the range [0%,100%] and is a normalized version of the former, which is the absolute position in terms of number of amino acids.

Next we compared the single- and di-nucleotide features. The overall pattern of the importance of single-nucleotides (Supplementary Figure 4) looks very similar to that from ref. ^11^, although without any feature directionality (favor/disfavored) for reasons described earlier in this section. To get a sense of the feature directionality, we used the L1-regularized regression model, and compared the weights there, to those found in ref. ^11^ (Supplementary Figure 8). Additionally, we do a paired plot of the actual predictions from the L1-regularized model and the gradient-boosted regression trees (Supplementary Figure 10), since we are here using the L1-regularized model to help understand the directionality of features. We see that overall, they are not radically different, and yet, the predictions diverge.

The dinucleotide features (Supplementary Figures 5-7) reveal numerous regions along the guide that show no evidence of single-nucleotide features, but strong evidence of dinucleotide importance. This is not surprising, given that removal of the dinucleotide features decreases the performance in a statistically significant manner (Supplementary Figure 11).

### Effect of different functional assays

Across all of our experiments, we observed that predictive performance was always better on the FC data as compared to the RES data. Given that the assay for functional knockdown was quite different for these data sets (FC used flow cytometry sorting, whereas RES used drug resistance), it would not be surprising to find that one of them had “cleaner” supervisory signal for machine learning. Another possibility is that the class of genes used in each is inherently easier or harder to predict on, but this explanation seems less likely. In any case, as a consequence of these observations of easier and harder data sets, we became interested in learning how these data sets interact with each other. In particular, we wanted to investigate how well each of them (FC, RES, FC+RES) served to generalize for each other. Thus, for all combinations of these data groups, we used one of them to train, and the other to test (Figure 5). We see that no matter what data was used to train, the RES data always had the worst performance. We also see that no matter what we tested, that using all of the available data (FC+RES) always had the maximal (and often the single best) performance. It is also interesting to recall that only for the FC data did addition of our new features not help the predictive performance, perhaps because this data is “easier,” and does not need as much help.

**Figure 5.**
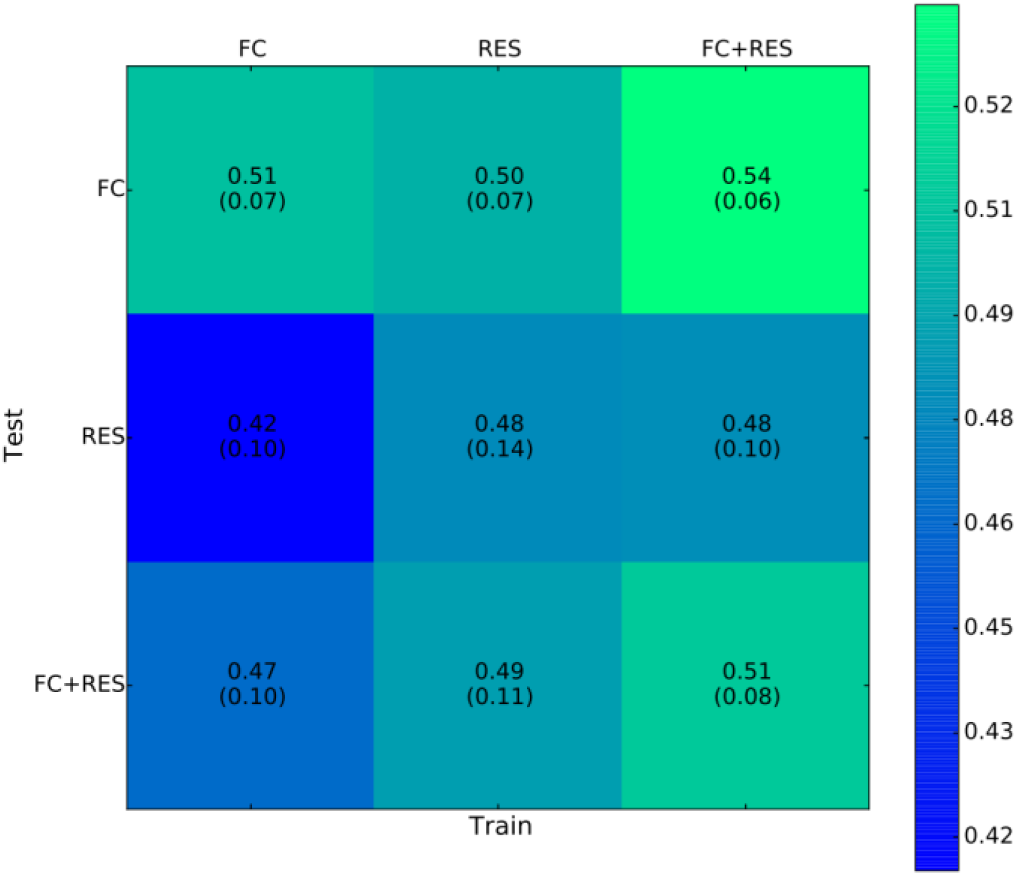
Spearman correlation when using different subsets of the data to train and test. The mean correlation across genes is written in each box, with the standard deviation in parenthesis below. Note that all experiments here used the Boosted Regression tree, with original+new features.

### Sensitivity to amount of training data

One desideratum when building predictive models, often hard to achieve, especially in biology, is to have enough data such that acquisition of further data would not help any further. That is, that the amount of training data available has in some sense saturated the model. To get a sense of how close we might be to such a setting, we systematically removed one gene further from the training data so as to artificially reduce the sample size, to see if we would observe any apparent plateaus. A plateau (*i.e*., flattening out of performance) as we add more genes is suggestive of having saturated the model—that is, it suggests that adding further data is not likely to greatly improve performance.

We see that the FC data appears to plateau slightly more than RES, but that neither data set appears to have saturated the model (Figure 6). These observations are consistent with the notion that the FC data are “easier” and the RES data “harder,” in that if one data set is harder because, for example, the supervisory signal is noisier, then it stands to reason that it would benefit from more data to overcome such noise. A similar argument can be made if the reason it is “harder” is not the noise, but some other property of the data.

**Figure 6.**
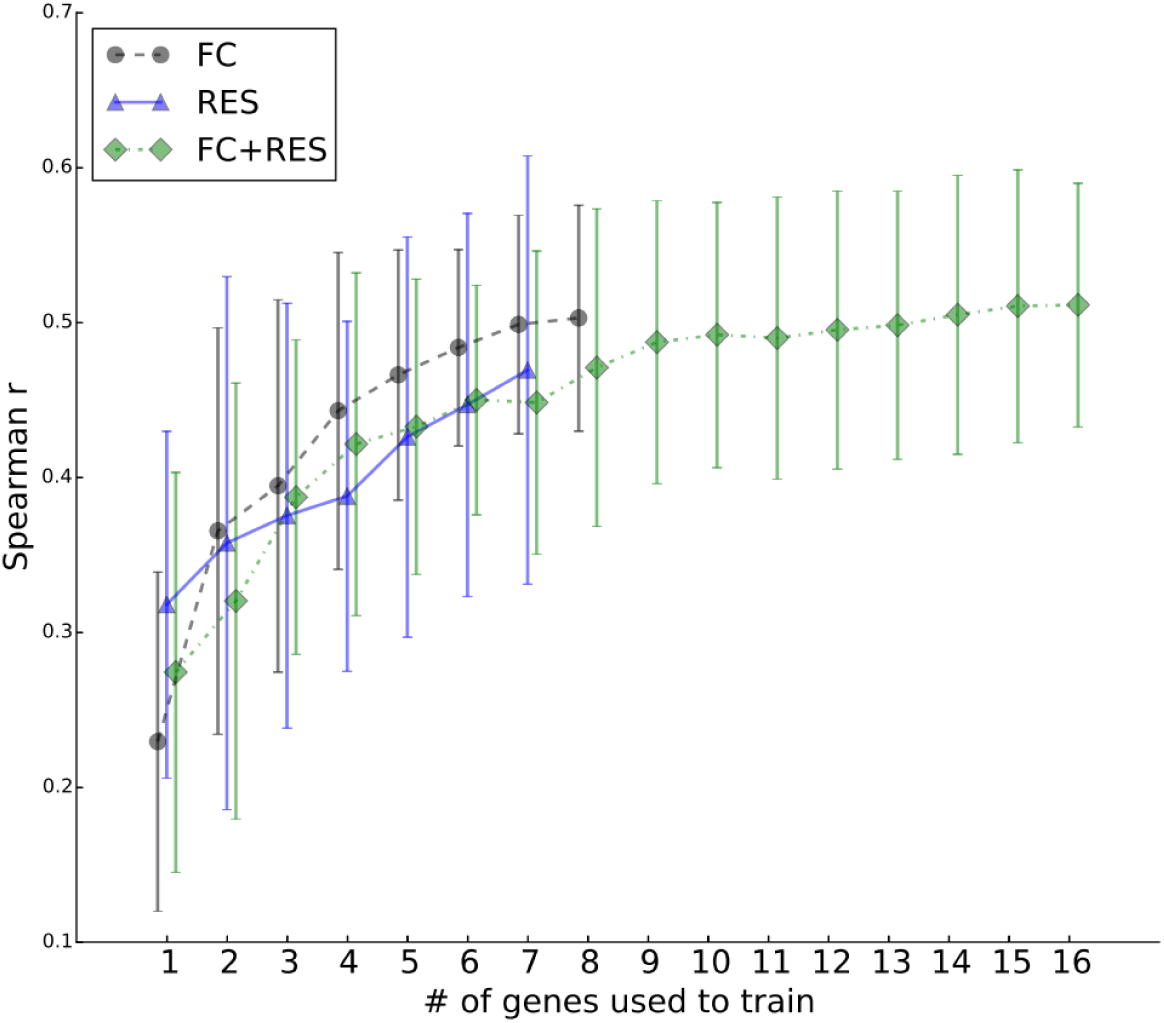
Investigation of the predictive performance as increasing numbers of genes (and guides) are available for training, using gradient-boosted regression trees with original+new features.

A plateau can be a marker of a model with too-limited capacity (too “simple” of a model), rather than sufficient data for the best possible model. As data set size increases, increasingly richer, higher capacity models^32^ should also be investigated. Note that the FC+RES comes much closer to a real plateau than either FC or RES alone, which is unsurprising as it contains roughly twice as much data as each individual data set.

## DISCUSSION

We have investigated, along several axes, how best to achieve optimal predicative performance for CRISPR/Cas9 gene editing, using a comprehensive set of guides over 17 genes for which we had a proxy to the knockout efficiency. The first and perhaps most important conclusion drawn was to avoid binarizing the data into active and inactive guides, instead retaining the real-valued normalized ranks for learning. Second, across a variety of statistical models, we found that Gradient-boosted regression trees yielded the best performance, and that addition of features not previously used, including thermodynamic features, PAM nucleotide interactions features, and position-independent nucleotide features yielded still further improvements. Inclusion of microhomology features, as well as some gene-specific features, did not yield noteworthy improvements in performance. Further, critical to any investigation of predictive models is the choice of an error measure. We have argued that for the particular task at hand, the Spearman correlation between the model predictions and actual normalized rank guide scores is the most appropriate choice, and that the AUC misses important information.

Although we did not report it in the main text, it is interesting to note that if one uses a stratified (across genes) cross-validation scheme, rather than a leave-one-gene-out cross-validation, performance increases. This suggests that there are gene-specific characteristics not presently well captured by our features, and that if one could find them, they would improve performance on the leave-one-gene out performance. In particular, such features would bridge the gap between the stratified and leave-one-gene out performances. In other words, imagine that if performance of a guide depended on whether its target gene had some combination of gene properties X, Y and Z. Then, in the absence of knowing X, Y and Z, we instead had training examples from the gene of interest (for example such as in cross-validation stratified by gene), we would still have good predictive performance. However, instead of having training examples from that gene of interest, it would suffice to have many examples covering the space of variables X, Y and Z, in order to learn the dependencies between them and other features for active guides. This then requires knowing what X, Y and Z are (should they exist). And these then would be new features that would bridge the gap between gene-stratified and leave-one-gene out cross-validation.

Further directions for investigation that might yield better predictive performance include the use of robust error losses, which can diminish the effect of outliers on the learned model. A brief investigation of the Huber loss for this purpose did not appear to help, but more work in this direction could be beneficial. The use of semi-supervised learning, enabling the inclusion of more guide/gene pairs than we have tested for efficacy would also be an interesting direction to pursue. In particular, the use of Deep Neural Networks in this setting could prove quite interesting. Additional features such as those based on string mismatch kernels^33^, gapped kmers^34^ and structural information of the guide/gene^18^ could also prove useful. With more data, new features may become relevant. For example, higher order nucleotide features (*e.g.*, order 3 nucleotides), for which we do not yet have enough data, could suddenly become useful. Another interesting area of investigation is the objective function used for parameter estimation. For the Gradient-boosted regression model the parameters were learned by minimizing the squared error between the true rank and the predicted rank, while for other models like logistic and linear regression, the (regularized) likelihood was maximized. However, it could be useful to investigate the use of maximizing the Spearman correlation itself, for example. Finally, in the gradient-boosted regression model, an extra set of parameters can be optimized—those controlling the maximum tree-depth, number of trees and the learning rate, for example, by way of Bayesian optimization.^35^

We investigated only on-target predictions, whereas in reality, for some applications, prediction of off-target effects are just as important. We plan in future work to model off-target predictions and then combine them with our on-target predictions for best overall performance, depending on the user’s needs.

## METHODS

### Data

A full description of the data generation can be found in ref. ^11^ and (Doench *et al*., in submission) and a summary of these data has already been provided at the start of the Results section. Here we describe details of how we computed the “guide scores” representing the effectiveness of any particular guide in knocking down its target gene. For each guide targeting a given gene, the log-fold-change (LFC) in abundance, as judged from read-counts was computed, where the change was with respect to whether the gene was knocked out or not (data from ref. ^11^ used flow cytometry, while Doench *et al*. used drug resistance—if knockout was successful, then the cell was resistant to the drug and survived). Next, normalized ranks were obtained for guides within each gene (or within each gene/drug pair if available) by assigning (possibly tied) ranks to each guide, and then re-scaling them to lie between 0, and 1. If more than one cell type was available, these normalized ranks were averaged across cell types just as in ref. ^11^ Thus, a final “guide score” for a given guide and gene (or gene-drug) was in [0,1], where 1 was indicative of successful knockdown, while 0 was unsuccessful knockdown. Note that normalized ranks were used instead of raw LFC because the LFC has a different maximum for genes that have more potential guides, suggesting that the LFC across different genes would not be meaningful. All processed data will be made available.

### Predictive Models

We used the following statistical models in our experiments: (1) linear regression, (2) L1-regularized linear regression, (3) L2-regularized linear regression, (4) the hybrid SVM plus logistic regression approach used in ref. ^11^, (5) Random Forest, (6) Gradient-boosted regression tree, (7) L1 logistic regression (a classifier), (8) SVM Classification. Implementations for each of these used scikit-learn package in python.^36^ For (2) and (3), we set the regularization parameter range to search over to be 100 points evenly spaced in log space, with a minimum of 10^-6^ and a maximum of 1.5 × 10^5^. Random Forest used the default setting, as did the Gradient-boosted regression trees (learning rate of 0.1, and 100 base estimators each with a maximum depth of three). For the SVM, we used a linear kernel with default L2 regularization unless otherwise noted (e.g. when reproducing ref. ^11^).

### Featurization

A “one-hot” encoding of the nucleotide sequences refers to taking a single categorical variable and converting it to more variables each of which can take on the value 0, or 1, with at most, one of them being “hot”, or on. For example, position 1 of the 30-mer guide plus context can take on A/C/T/G, and it gets converted to four binary variables, one for each possible nucleotide. These are “order 1” features. For “order 2” features, we looked at all adjacent pairwise nucleotides as features, such as AA/AT/AG/etc. There are 4 × 4 = 16 such pairs, thus a single variable representing one such pair gets one-hot encoded into 16 binary variables. In ref. ^11^, only “position-specific” nucleotide features were used in this manner, meaning that for each position on the guide, a different one-hot-encoded feature was used. Here, however, taking inspiration from some of the string kernel literature^37^, we augmented these features sets by also including “position-independent” features, where, for order 1 features for example, this would mean a feature for how many A’s and how many T’s, etc. were in the guide, ignoring their position in the guide, and similarly for “order 2”. Therefore, for a 30mer (20mer guide plus context), we obtained 80 order 1 and 320 order 2 position-specific features, and 4 order 1 and 16 order 2 position-independent features.

GC counts features were computed as in ref. ^11^, which is to say the number of Gs and Cs in the 30mer were counted, yielding one feature, and then another feature with count>10 was also used.

The two nucleotides in the N and N positions relative to the PAM “NGGN” were one-hot encoded yielding 16 features, one for each NN possibility (*e.g.*, AT).

Thermodynamic features were computed from the melting temperatures of the DNA version of the RNA guide sequence, or portions thereof, using the Biopython Tm_staluc function.^8,29^ In addition to using the melting temperature of the entire 30mer guide plus context, we also separated the thermodynamic features into three further features corresponding to the melting temperature of three distinct parts of the guide—in particular, the 5 nucleotides immediately proximal to the PAM, the 8 nucleotides adjacent to that (away from the PAM), and then the 5 nucleotides in turn adjacent to the 8mer (again, away from the PAM).

Guide positional features such as amino acid cut position and percent peptide were computed as in ref. ^11^ and correspond to how far from the start of the protein coding region of the gene the guide was positioned.

Not all features will appear in our Supplementary files, as features that did not vary at all were removed from all computations.

### Stratification/CV

Other than otherwise noted (*i.e*., the section Sensitivity to amount of training data), cross-validation was always performed by leaving all guides for one gene out at a time. For the inner cross-validation in the required nested cross-validation used to set the regularization weights for L1/L2, we also used leave-one-gene-out cross-validation. In the section ‘Sensitivity to amount of training data’, we decreased the size of the training data set by systematically removing all guides for one gene, and we removed that gene each time (*i.e*., each fold) at random.

When examining feature importance, we averaged either the Gini importance (for gradient boosted regression trees), or the weights (for L1-regularized regression) across genes.

### Statistical significance

The only test of statistical significance used was to compare the Spearman correlation prediction measure between two approaches when using exactly the same test data and cross-validation. To do so, we used three sets of numbers: (i) the Spearman correlation performance measure for one approach, (ii) the Spearman correlation performance measure for the other approach, and (iii) the Spearman correlation between the predictions from each approach. These values were used in conjunction with a method to test dependent measures of correlations (here they are dependent because each approach uses the same ground truth).^40^ The precision of the implementation for this method resulted in the occasional p-value of 0.0. In such cases, we instead report p<1 × 10^-16^, the smallest p-value that we observed the method reporting (strictly speaking this depends on the number of samples, and correlation between predictions, but it seemed a reasonably practical place-holder). Also of note is that this measure of statistical significance does not give equal weight to each gene, rather, each gene is represented in proportion to its number of guides, unlike the bar plot figures we have used throughout.

### Software and Code availability

All of our source code, experiments, data, figure-generation code, and a public cloud-based prediction server will be made available from http://research.microsoft.com/en-us/projects/azimuth.

## Supplementary Materials

### Supplementary Figures

**Supplementary Figure 1.**
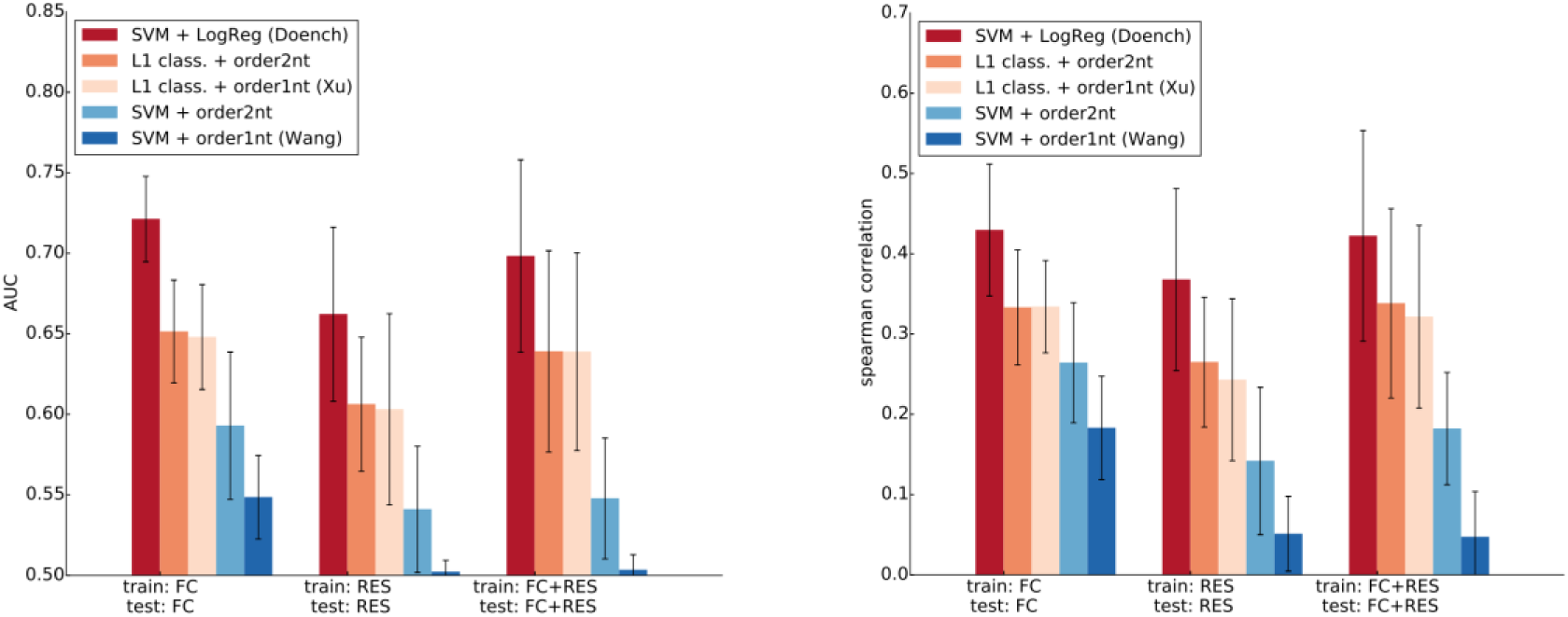
Comparison of the classification methods, including that from ref. ^11^ (SVM + LogReg), a baseline L1 classifier (using order 1 and 2 features), an L1classifier with only order 1 features (*i.e.,* the modeling approach of Xu *et al* ^16^), the classification method from ref. ^4^ which uses only “order 1” nucleotide features and nothing else (Wang), a version of Wang with both “order 1” and “order 2” features. For completeness, we show both AUC and Spearman correlation predictive performance. Even when using the modeling approaches of others (Wang, Xu), we always trained and tested on the same data, as noted in the horizontal labels. The statistical significances of the improvement in Spearman correlations given by the SVM+LogReg model, when comparing to, respectively: L1 class. (ord. 2), L1 class. (ord. 1), Wang + features, Wang, are *p* = 1.8 × 10^-8^, *p* = 3.2 × 10^-5^, *p* = 1.3 × 10^-14^, *p* < 10^-16^ (FC), *p* = 5.2 × 10^-13^, *p* < 10^-16^, *p* < 10^-16^, *p* < 10^-16^ (RES), *p* < 10^-16^, *p* < 10^-16^, *p* < 10^-16^, *p* < 10^-16^ (FC+RES). The statistical significance of the improvement in Spearman correlation of the L1 classifier with order 2 (which includes 1), compared to only using order 1 is *p* = 0.03, *p* = 1.8 × 10^-4^, *p* = 1.1 × 10^-6^, showing that the order 2 features do help, although less so than with the gradient-boosted regression trees (Supplementary Figure 10).

**Supplementary Figure 2.**
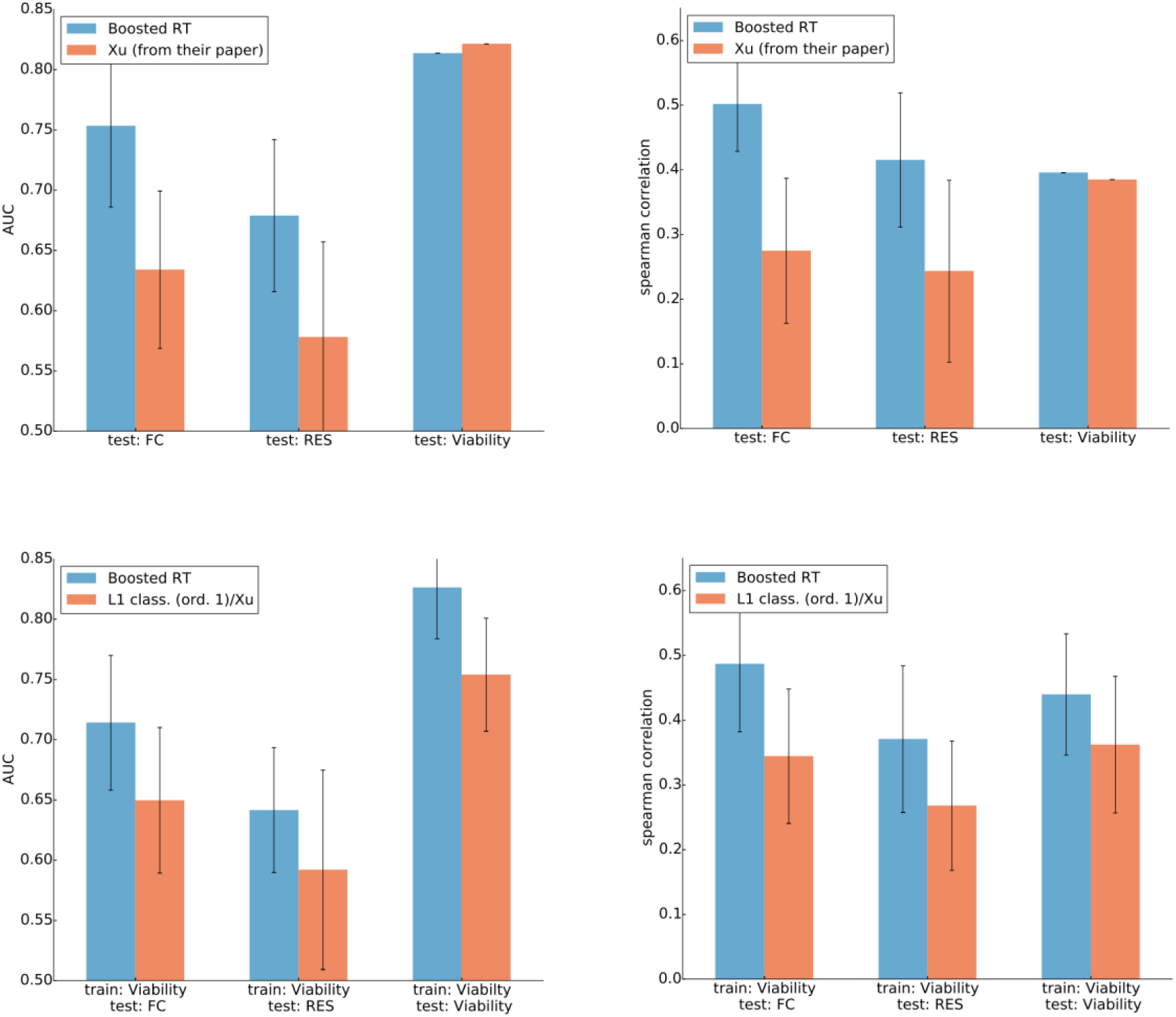
Top row: comparison of our chosen modeling approach (boosted regression trees, “Boosted RT”), against the final, trained model provided by Xu *et al* ^16^. The horizontal axis denotes which set of data was used to test on—the FC data in its entirety, the RES data set in its entirety, the Viability data from Xu *et al.* Bottom row: in contrast to the top row, here we re-trained the Xu *et al.* model (L1 classification, order 1 nuc. features) on the Viability data, and then tested it on FC, RES, and Viability data, but now using stratified (by gene) cross-validation in all cases, so that the training and test sets were never overlapping.

**Supplementary Figure 3.**
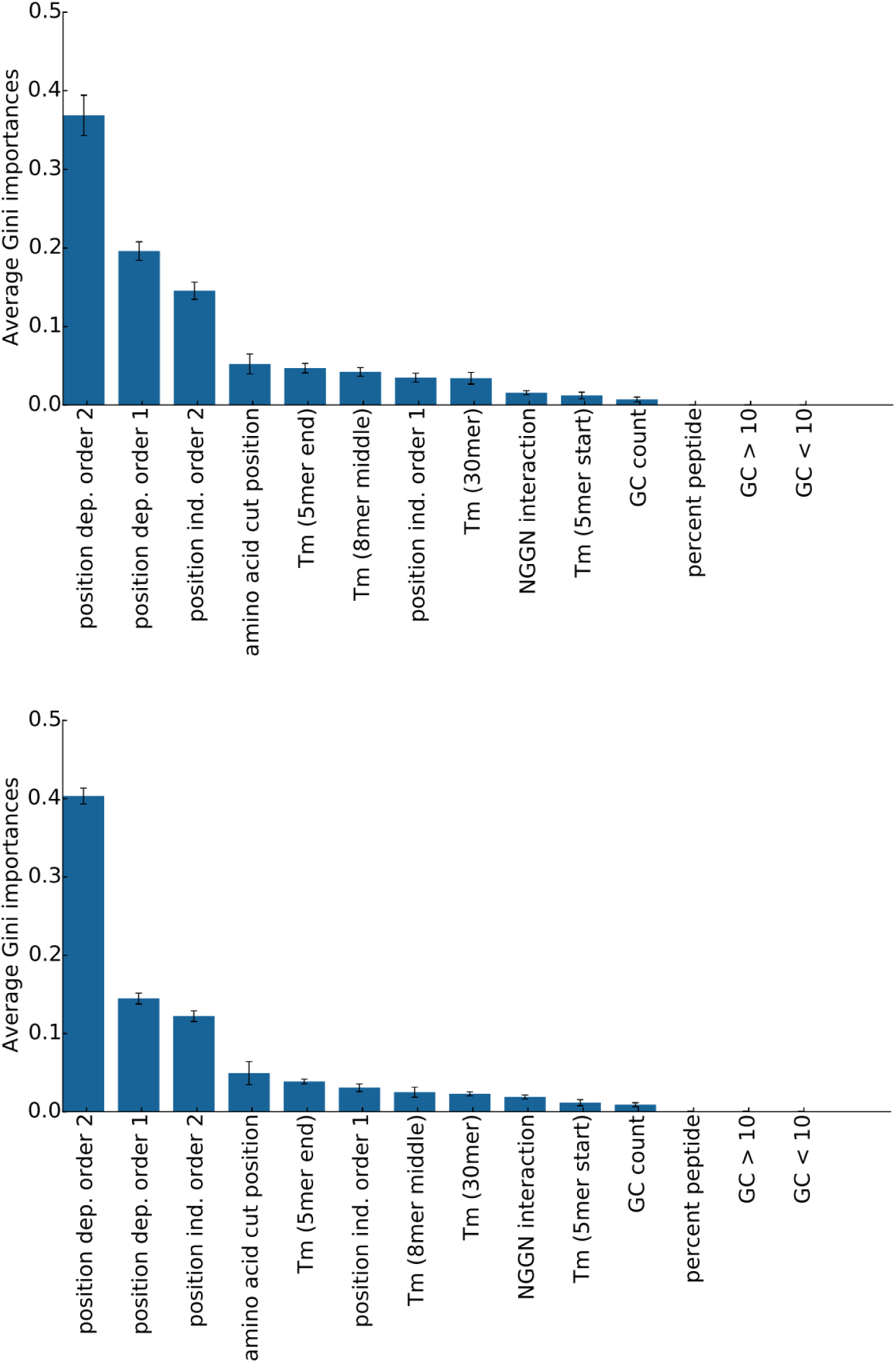
Relative importance of each group of features in the Gradient-boosted regression trees using the Gini importance, for the experiment using the FC data and (top) the RES data (bottom). FC+RES is Figure 4 in the main manuscript. “position dep. order 2” denotes position-dependent features for adjacent paired nucleotide features, whereas “position ind. order 2” denotes the same thing, but aggregated over all positions.

**Supplementary Figure 4.**
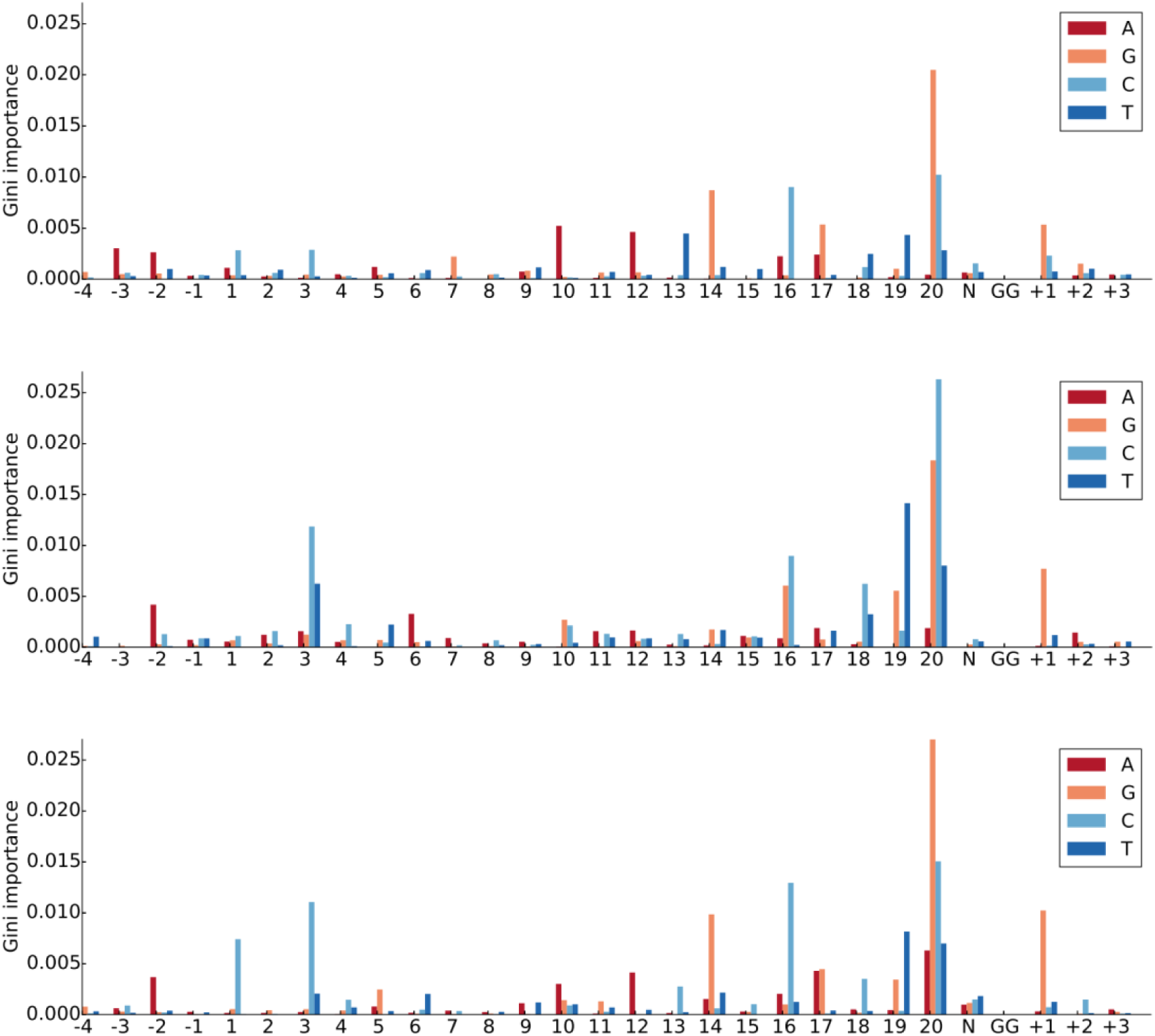
Gini importance of “order 1” nucleotide features in the Boosted regression trees, for each of the FC (top), RES (middle) and FC+RES (bottom) data sets.

**Supplementary Figure 5.**
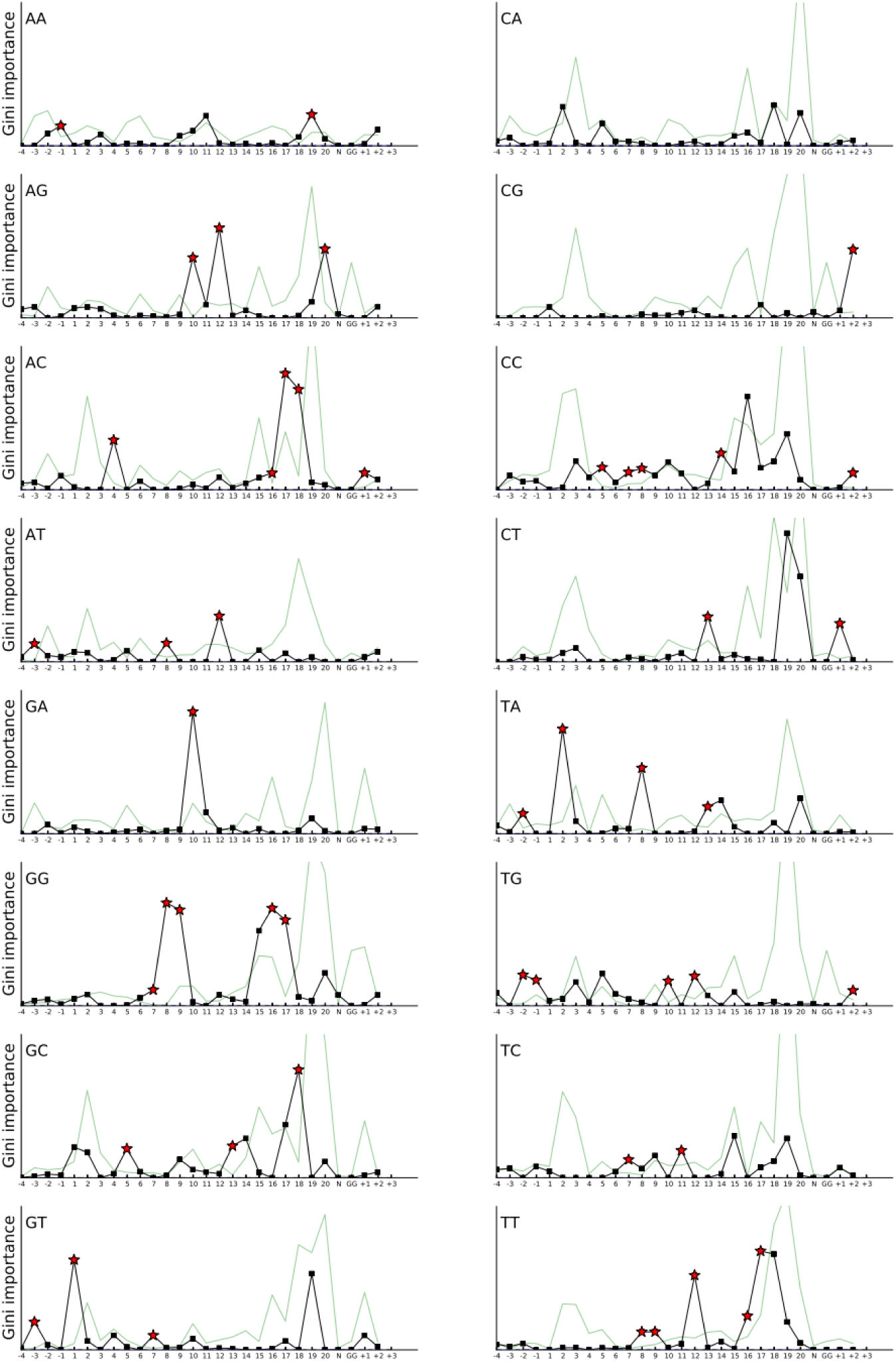
Gini importance of dinucleotide features in the Boosted regression trees for the FC data (black). The green line shows the average Gini importance for each of the position-dependent single nucleotide features that correspond to the given dinucleotide. Therefore, when the black line is higher than the green line, the dinucleotide features more “surprising”, as there was no hint of them in the single nucleotide features. Those that were most surprising (more black line more than double the green line, and with at least a value of 1e-3), are denoted with a red star. When the vertical axis would have cut off the red-star, the red star is plotted at the top of the axis. The vertical axis is the same in each axis, with a maximum value of Gini importance of 0.01.

**Supplementary Figure 6.**
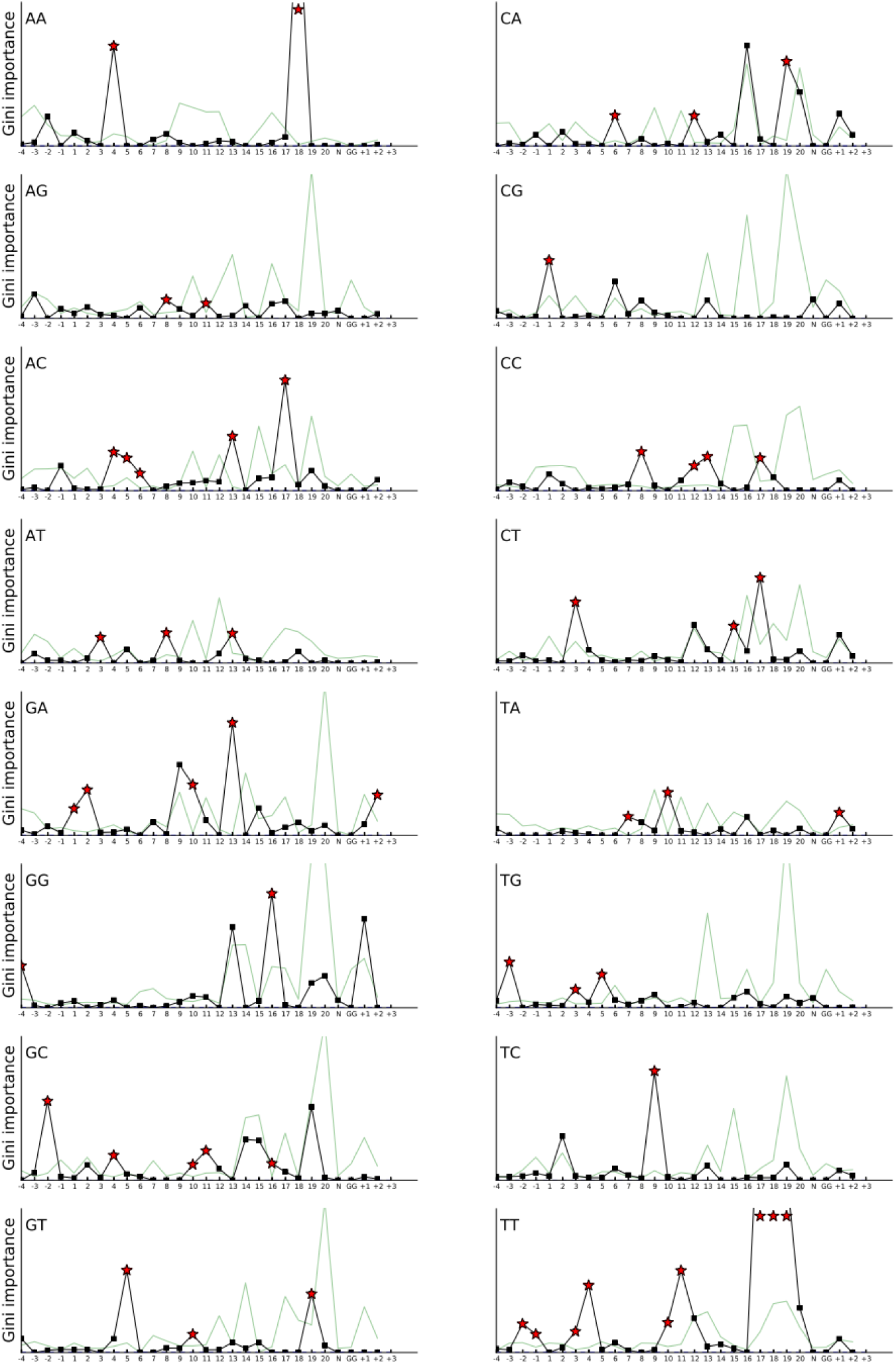
Gini importance for dinucleotide features for the RES data set. See Supplementary Figure 5 for more information.

**Supplementary Figure 7.**
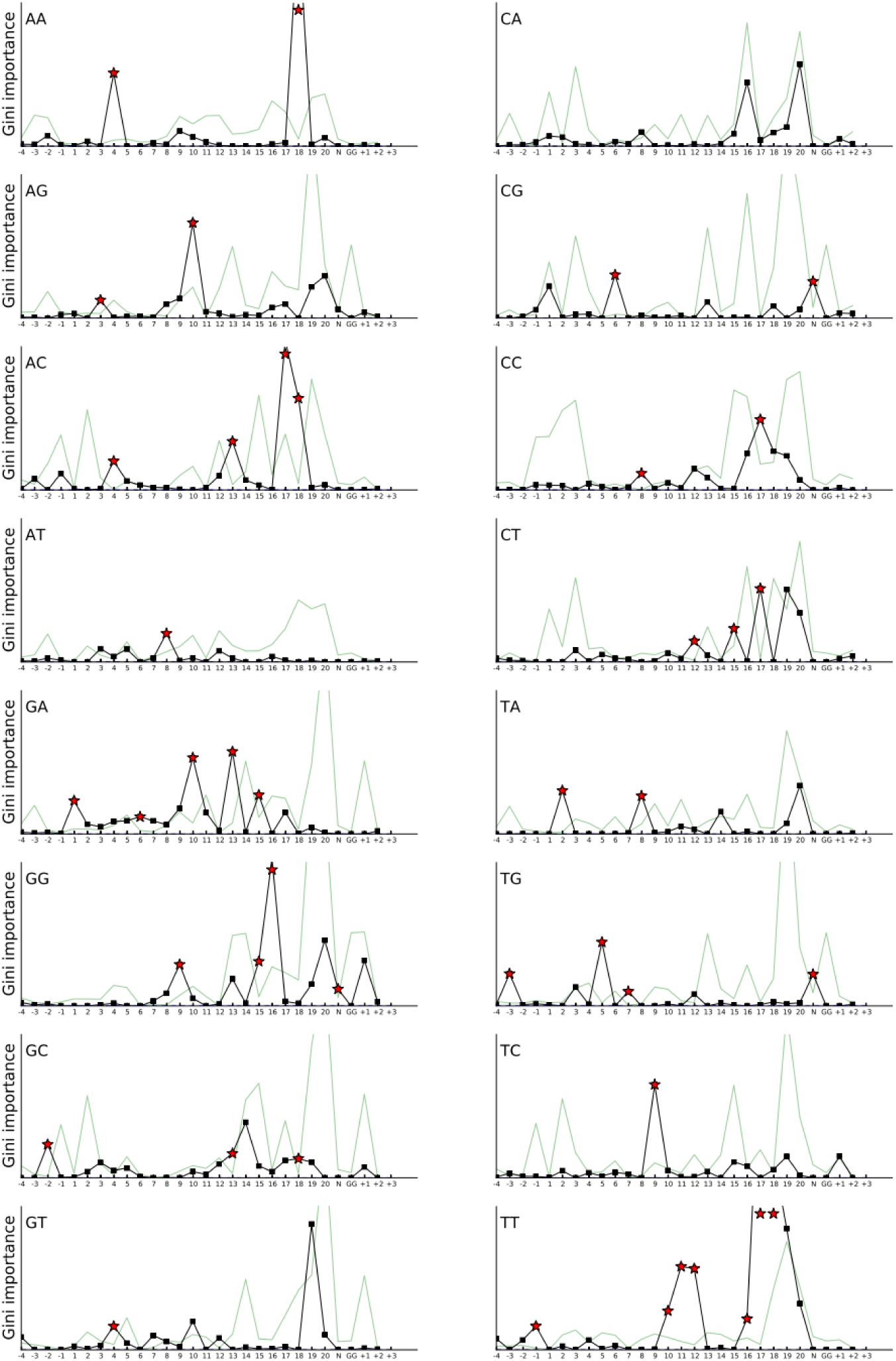
Gini importance for dinucleotide features for the RES data set. See Supplementary Figure 5 for more information.

**Supplementary Figure 8.**
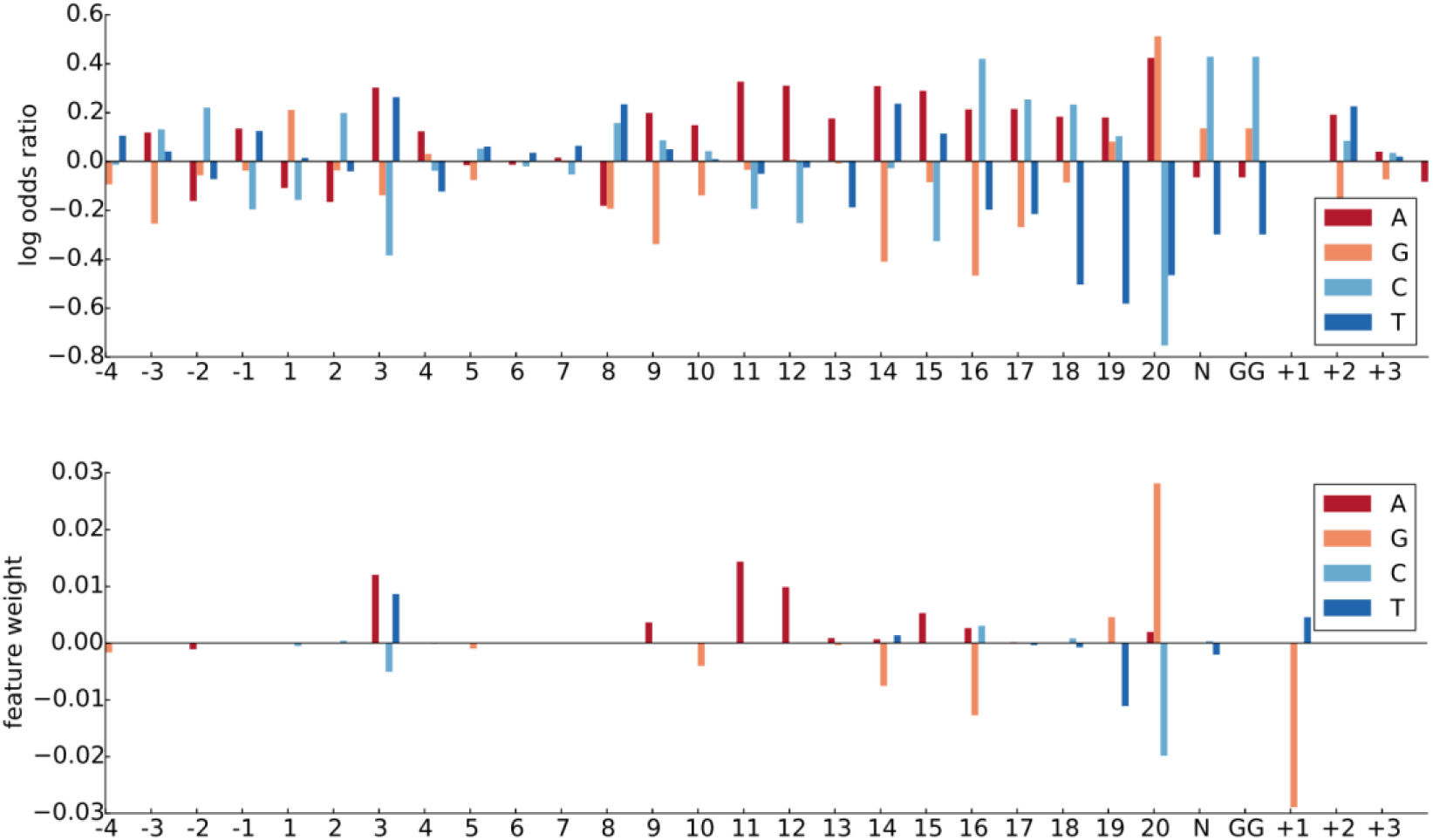
Comparison of single nucleotide feature weights (i.e, log odds ratio) from ref. ^11^ (top) to the weights from our L1-regularized regression, on the FC data, using the original features only (bottom). Note because the top model is a classification model, whereas the bottom plot is a regression model, the vertical axis is not comparable between the two even though each represents the feature weight.

**Supplementary Figure 9.**
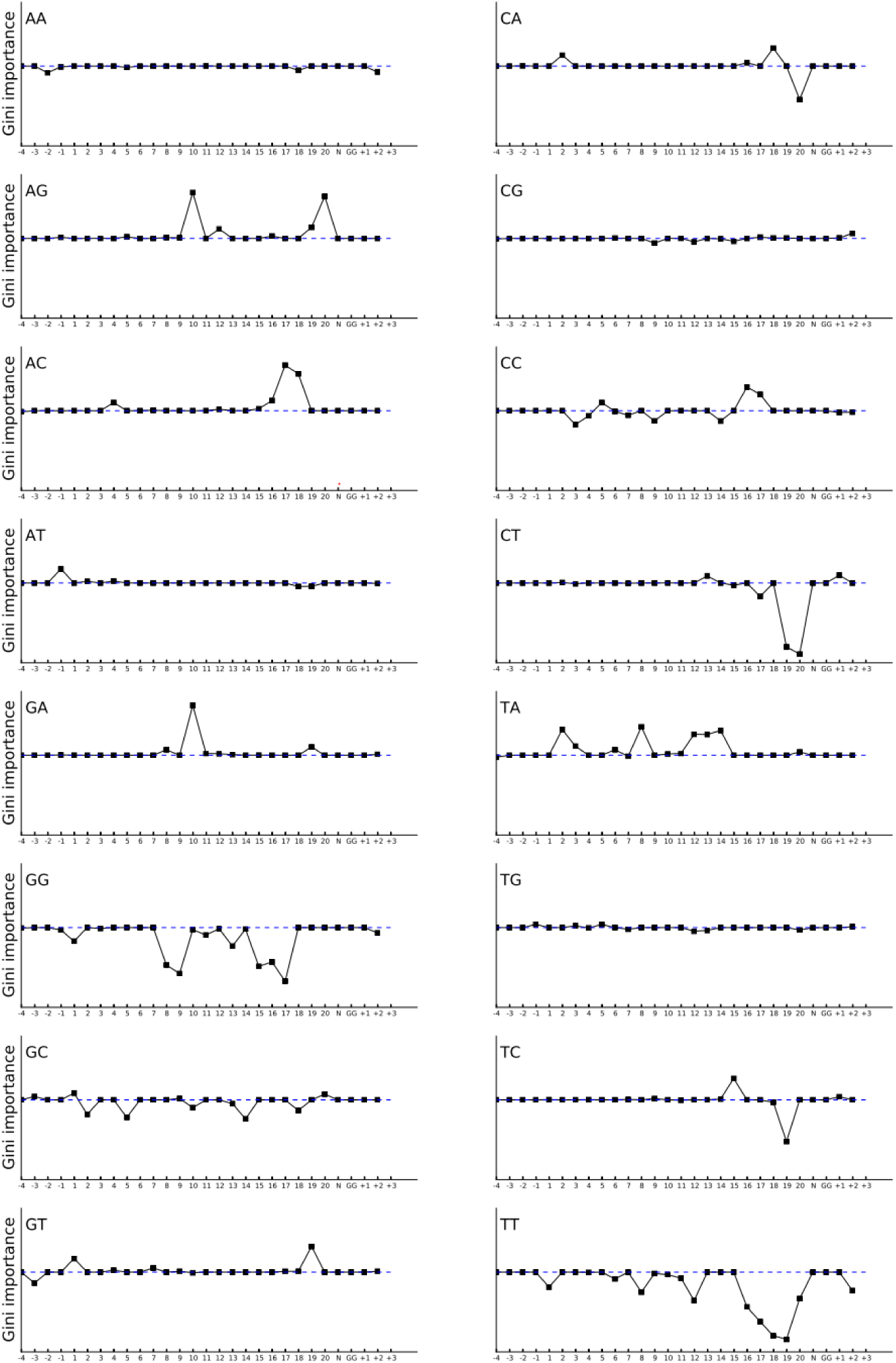
The L1-regularized weights denoting feature importance, for the dinucleotide features in the FC data (black). Dashed blue line denotes zero weight. The vertical axis is the same in each axis, with a minimum value of -0.025 and maximum of 0.02.

**Supplementary Figure 10.**
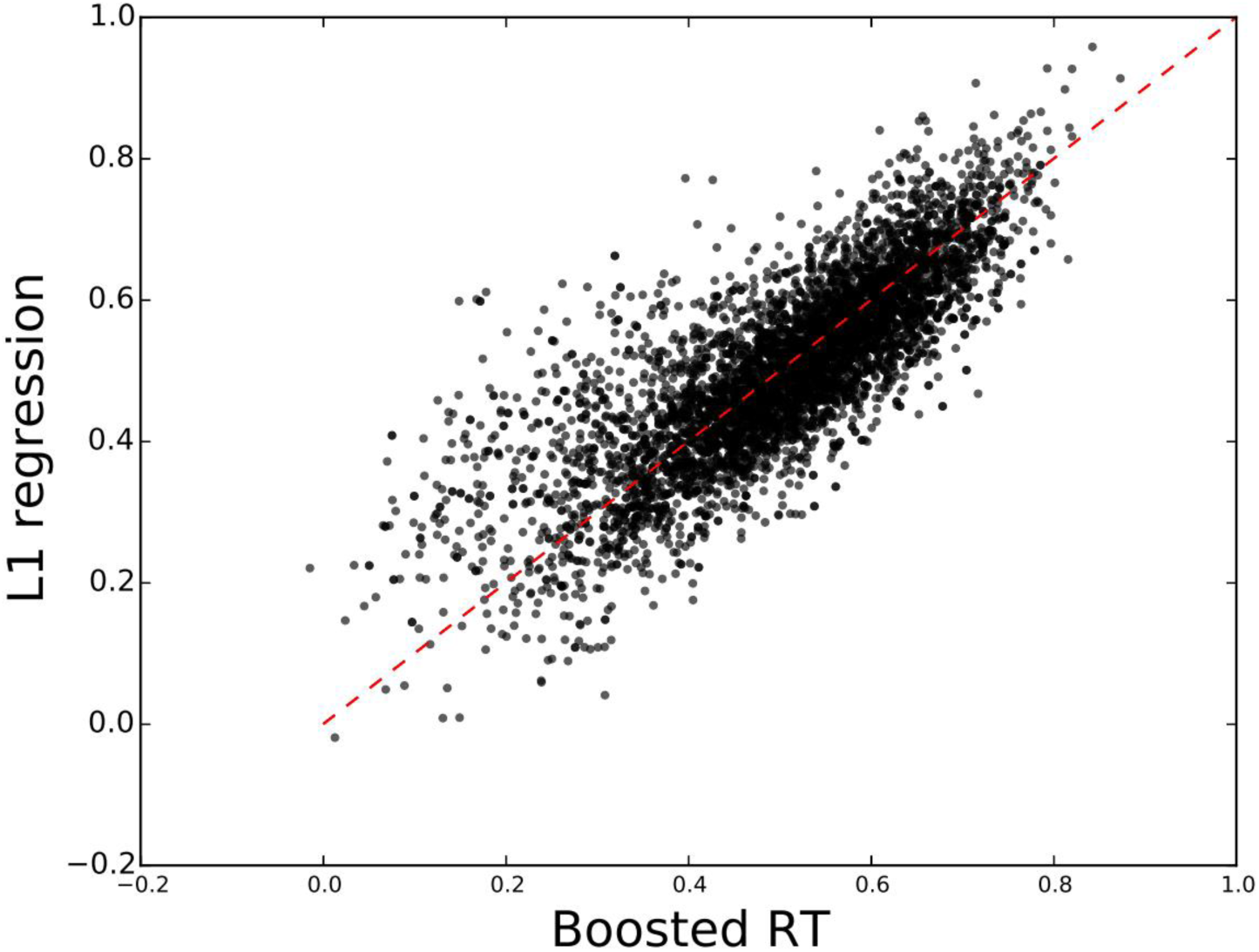
Paired plot of the model predictions from gradient-boosted regression trees and L1-regularized regression, on the FC+RES data using original plus new features. Dashed redlines denotes the diagonal. Note that model predictions can on occasion become smaller than zero, as the model does not constrain them otherwise.

**Supplementary Figure 11.**
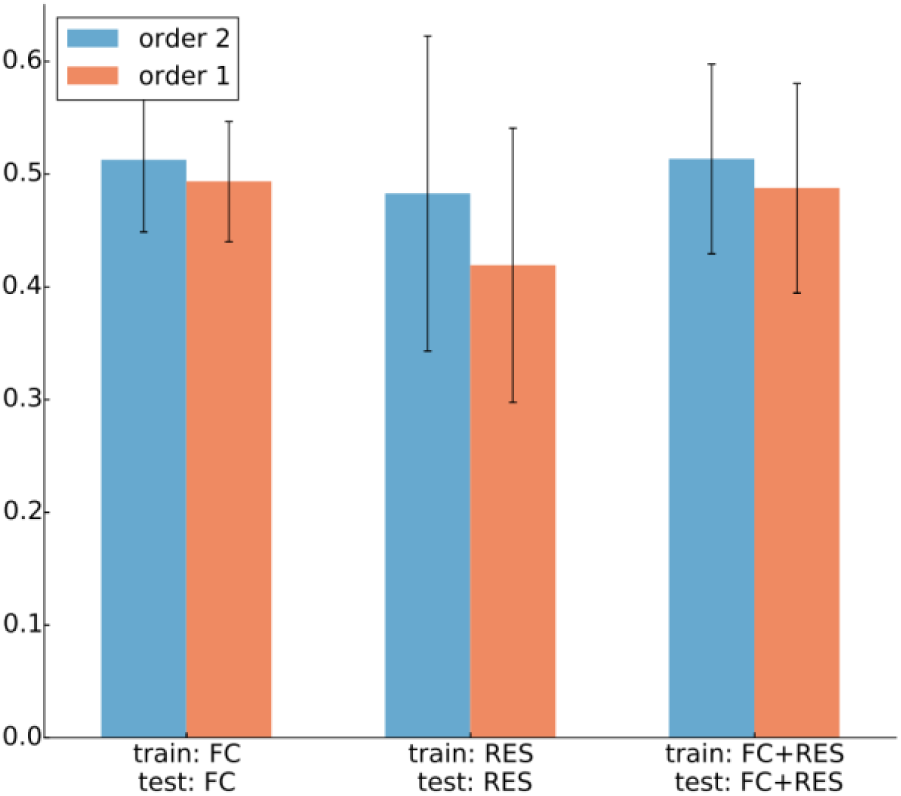
Comparison of Gradient Boosted regression trees with our best feature set (which includes both order 1 and order 2 features), and then again, removing the order 2 features. The statistical significance between the Spearman correlation over all genes between the two sets of features, for each data set from left to right is, respectively, *p* = 4 × 10^-4^, *p* = 8 × 10^-9^, *p* = 2 × 10^-11^.

Links to Supplementary Information files (to be released):

1. feature_importances, boosted regression tree, all features, train FC, test FC
2. feature_importances, boosted regression tree, all features, train RES, test RES
3. feature_importances, boosted regression tree, all features, train FC+RES, test FC+RES
4. Processed FC+RES data

## References

1. Doudna, J. A. & Charpentier, E. The new frontier of genome engineering with CRISPR-Cas9. Science (80-.). 346, 1258096–1258096 (2014).

2. Gaj, T., Gersbach, C. A. & Barbas, C.F. Zfn, TALEN, and CRISPR/Cas-based methods for genome engineering. Trends Biotechnol. 31, 397–405 (2013).

3. Shalem, O. et al. Genome-Scale CRISPR-Cas9 Knockout Screening in Human Cells. Science (80-.). 343, 84–87 (2013).

4. Wang, T., Wei, J. J., Sabatini, D. M. & Lander, E. S. Genetic screens in human cells using the CRISPR-Cas9 system. Science 343, 80–4 (2014).

5. Koike-Yusa, H., Li, Y., Tan, E.-P., Velasco-Herrera, M. D. C. & Yusa, K. Genome-wide recessive genetic screening in mammalian cells with a lentiviral CRISPR-guide RNA library. Nat. Biotechnol. 32, 267–73 (2014).

6. Mandal, P. K. et al. Efficient Ablation of Genes in Human Hematopoietic Stem and Effector Cells using CRISPR/Cas9. Cell Stem Cell 15, 643–652 (2014).

7. Schwank, G. et al. Functional repair of CFTR by CRISPR/Cas9 in intestinal stem cell organoids of cystic fibrosis patients. Cell Stem Cell 13, 653–8 (2013).

8. Shen, H. Precision gene editing paves way for transgenic monkeys. Nature 503, 14–5 (2013).

9. Long, C. et al. Prevention of muscular dystrophy in mice by CRISPR/Cas9-mediated editing of germline DNA. Science 345, 1184–1188 (2014).

10. Hsu, P. D. et al. DNA targeting specificity of RNA-guided Cas9 nucleases. Nat. Biotechnol. 31, 827–32 (2013).

11. Doench, J. G. et al. Rational design of highly active sgRNAs for CRISPR-Cas9-mediated gene inactivation. Nat Biotechnol (2014). doi:10.1038/nbt.3026

12. Hartenian, E. & Doench, J. G. Genetic screens and functional genomics using CRISPR/Cas9 technology. FEBS J. 282, 1383–93 (2015).

13. Stemmer, M., Thumberger, T., del Sol Keyer, M., Wittbrodt J., & Mateo, J. L. CCTop: An Intuitive, Flexible and Reliable CRISPR/Cas9 Target Prediction Tool. PLoS One 10, e0124633 (2015).

14. Heigwer, F., Kerr, G. & Boutros, M. E-CRISP: fast CRISPR target site identification. Nat. Methods 11, 122–123 (2014).

15. Montague, T. G., Cruz, J. M., Gagnon, J. a., Church, G. M. & Valen, E. CHOPCHOP: A CRISPR/Cas9 and TALEN web tool for genome editing. Nucleic Acids Res. 42, 401–407 (2014).

16. Xu, H. et al. Sequence determinants of improved CRISPR sgRNA design. Genome Res. (2015). doi:10.1101/gr.191452.115

17. Ghosal, S., Das, S. & Chakrabarti, J. Computational Approaches for Designing Efficient and Specific siRNAs. (2012).

18. Li, W. & Cha, L. Genetic studies of diseases: Predicting siRNA efficiency. Cell. Mol. Life Sci. 64, 1785–1792 (2007).

19. Knott, S. R. V et al. A Computational Algorithm to Predict shRNA Potency. 1–12 (2014).

20. Breiman, L., Friedman, J., Olshen, R. & Stone, C. Classification and Regression Trees. (Wadsworth and Brooks, 1984).

21. Järvelin, K. & Kekäläinen, J. Cumulated Gain-based Evaluation of IR Techniques. ACM Trans. Inf. Syst. 20, 422–446 (2002).

22. Anders, C., Niewoehner, O., Duerst, A. & Jinek, M. Structural basis of PAM-dependent target DNA recognition by the Cas9 endonuclease. Nature (2014). doi:10.1038/nature13579

23. Wu, X. et al. Genome-wide binding of the CRISPR endonuclease Cas9 in mammalian cells. Nat. Biotechnol. 32, 670–676 (2014).

24. Bae, S., Kweon, J., Kim, H. S. & Kim, J.-S. Microhomology-based choice of Cas9 nuclease target sites. Nat. Methods 11, 705–706 (2014).

25. Freund, Y. & Schapire, R. E. A Decision-theoretic Generalization of On-line Learning and an Application to Boosting. in Proc. Second Eur. Conf. Comput. Learn. Theory 23–37 (Springer-Verlag, 1995). at <http://dl.acm.org/citation.cfm?id=646943.712093>

26. Schapire, R. E. The Strength of Weak Learnability. Mach. Learn. 5, 197–227 (1990).

27. Review, P. A., Random, G. & Tech-, C. F. C. E. Rasmussen & C. K. I. Williams, Gaussian Processes for Machine Learning, the MIT Press, 2006, ISBN 026218253X. c2006 Massachusetts Institute of Technology. www.GaussianProcess.org/gpml. (2006).

28. Hinton, G. & Salakhutdinov, R. Reducing the Dimensionality of Data with Neural Networks. Science (80-.). 313, 504–507 (2006).

29. Bengio, Y. Learning Deep Architectures for AI. Found. Trends(r) Mach. Learn. 2, 1–127 (2009).

30. Zou, H. & Hastie, T. Regularization and variable selection via the Elastic Net. J. R. Stat. Soc. Ser. B 67, 301–320 (2005).

31. Menze, B. H. et al. A Comparison of Random Forest and its Gini Importance with Standard Chemometric Methods for the Feature Selection and Classification of Spectral Data. BMC Bioinformatics 10:213, (2009).

32. Thodberg, H. H. Improving generalization of neural networks through pruning. Int. J. Neural Syst. 1, 317–326 (1991).

33. Leslie, C. & Noble, W. S. Mismatch String Kernels for SVM Protein. Text

34. Ghandi, M., Lee, D., Mohammad-Noori, M. & Beer, M. A. Enhanced regulatory sequence prediction using gapped k-mer features. PLoS Comput. Biol. 10, e1003711 (2014).

35. Mockus, J. B. & Mockus, L. J. Bayesian approach to global optimization and application to multiobjective and constrained problems. J. Optim. Theory Appl. 70, 157–172 (1991).

36. Pedregosa, F. et al. Scikit-learn: Machine Learning in {P}ython. J. Mach. Learn. Res. 12, 2825– 2830 (2011).

37. Ben-Hur, A., Ong, C. S., Sonnenburg, S., Schölkopf, B. & Rätsch, G. Support vector machines and kernels for computational biology. PLoS Comput. Biol. 4, (2008).

38. Le Novere, N. MELTING, computing the melting temperature of nucleic acid duplex. Bioinformatics 17, 1226–1227 (2001).

39. Cock, P. J. A. et al. Biopython: freely available Python tools for computational molecular biology and bioinformatics. Bioinformatics 25, 1422–3 (2009).

40. Steiger, J. H. Tests for comparing elements of a correlation matrix. Psychol. Bull. 87, 245–251 (1980).

